# Expanded huntingtin CAG repeats disrupt the balance between neural progenitor expansion and differentiation in human cerebral organoids

**DOI:** 10.1101/850586

**Authors:** Jinqiu Zhang, Jolene Ooi, Kagistia Hana Utami, Sarah R. Langley, Obed Akwasi Aning, Dong Shin Park, Magdalena Renner, Shiming Ma, Chit Fang Cheok, Juergen A. Knoblich, Florent Ginhoux, Enrico Petretto, Mahmoud A. Pouladi

## Abstract

Huntington disease (HD) manifests in both adult and juvenile forms. Mutant *HTT* gene carriers are thought to undergo normal brain development followed by a degenerative phase, resulting in progressive clinical manifestations. However, recent studies in children and prodromal individuals at risk for HD have raised the possibility of abnormal neurodevelopment. Although key findings in rodent models support this notion, direct evidence in the context of human physiology remains lacking. Using a panel of isogenic HD human embryonic pluripotent stem cells and cerebral organoids, we investigated the impact of mutant HTT on early neurodevelopment. We find that ventricular zone-like neuroepithelial progenitor layer expansion is blunted in an *HTT* CAG repeat length-dependent manner due to premature neurogenesis in HD cerebral organoids, driven by cell intrinsic processes. Transcriptional profiling and imaging analysis revealed impaired cell cycle regulatory processes, increased G1 length, and increased asymmetric division of apical progenitors, collectively contributing to premature neuronal differentiation. We demonstrate increased activity of the ATM-p53 pathway, an up-stream regulator of cell cycle processes, and show that treatment with ATM antagonists partially rescues the blunted neuroepithelial progenitor expansion in HD organoids. Our findings suggest that CAG repeat length regulates the balance between neural progenitor expansion and differentiation during early neurodevelopment. Our results further support the view that HD, at least in its early-onset forms, may not be a purely neurodegenerative disorder, and that abnormal neurodevelopment may be a component of HD pathophysiology.

## Introduction

Huntington disease (HD) is a fatal, autosomal dominant neurodegenerative disorder. The disease is caused by a polymorphic CAG repeat expansion, encoding an extended polyglutamine (polyQ) tract, in the huntingtin *(HTT)* gene to greater than 36 repeats [1]. There is an inverse relationship between the length of the pathogenic CAG repeat and rate of disease onset, with repeat lengths in the 40’s generally resulting in disease onset in the fourth decade whereas repeat lengths greater than 60 lead to juvenile onset in childhood (<10 years) or adolescence (<20 years) [1,2].

Given the age-dependent nature of the disorder, studies to date have largely focused on understanding the degenerative processes that manifest in adulthood. However, studies in prodromal individuals and children at risk for the disease have raised the possibility that mutant HTT may lead to abnormal neurodevelopment, in addition to its neurodegenerative effects in HD. For example, intracranial volume, a surrogate of maximal brain growth achieved during development, is smaller in pre-manifest HD mutation carriers [3]. Similarly, head circumference, which correlates closely with intracranial volume, is smaller in children who carry the HD mutation and who are estimated to be decades from onset [4]. Moreover, the morphology of brain regions is substantially different in adult pre-manifest HD mutation carriers, with increased volume of the cerebral cortex and decreased volume of the basal ganglia and cerebral white matter [5]. A recent longitudinal study in children and adolescent HD gene carriers who were estimated to be on average 35 years from clinical onset revealed that an initial hypertrophy of striatal volume precedes the volume decline that develops with age [6]. Furthermore, two recent whole-exome sequencing studies have identified compound heterozygous variants in the *HTT* gene in patients with Rett-like syndrome, a neurodevelopmental disorder [7,8]. These findings suggest that mutations in *HTT* can lead to abnormal neurodevelopment.

Studies in mouse models support this notion and have demonstrated a role for normal HTT in brain development. Loss of HTT has been shown to cause profound neurodevelopmental abnormalities, including striatal and cortical malformation [9–11]. In early corticogenesis, HTT was shown to be required for proper mitotic spindle orientation, a key determinant of cortical progenitor fate [12]. Loss of HTT in cortical progenitors decreased their proliferation [12,13] and promoted their differentiation [12]. Furthermore, depletion of HTT impaired the polarity and migration of post-mitotic neurons [14,15], processes that are critical for neuronal identity specification and cortical lamination [16,17].

Studies have also implicated mutant HTT in abnormal neurodevelopment. Similar to loss of normal HTT, mutant HTT has been shown to disrupt mitotic spindle orientation in neural stem cells [18, 19] and cortical progenitors [20], as well as the migration of post-mitotic neurons [14]. However, the impact of mutant HTT on early neurogenesis is less clear. Indeed, studies examining neurogenesis using mouse HD stem cells have given conflicting results showing that mutant HTT both impairs neurogenesis [21] and promotes it [13,22]. Efforts to examine the impact of mutant HTT on neurodevelopment and neurogenesis using human pluripotent stem cells (hPSCs) having been similarly unclear [18,19,23–26] (reviewed in [27]). Mutant HTT was shown to impact early ectodermal development, reflected in an altered phenotypic signature, in an in vitro model of neurulation [28]. Based on transcriptional analyses of hPSCs [26], neural progenitors [23,25], and neurons [24], a number of studies have also suggested that mutant HTT impairs neural progenitor differentiation and delays neuronal maturation. However, assessment of cortical neurogenesis showed no effect of mutant HTT on the quantity or timeline of differentiation of deep and upper layer cortical neurons [24]. These studies have largely relied on directed differentiation of embryonic or induced pluripotent stem cells in two-dimensional culture systems to induce neurogenesis. One exception is a recent study using tridimensional HD hPSC-derived cortical organoids which presented evidence of delayed neuronal maturity based on transcriptional profile analysis, suggesting that mutant HTT precludes normal neuronal fate acquisition [29].

Here, using a novel panel of isogenic HD human embryonic pluripotent stem cells and cerebral organoids, we investigated the impact of mutant HTT and CAG repeat length on early neurodevelopment. Cerebral organoid-based modeling of early neurodevelopment helps overcome some of the limitations of two-dimensional neuronal systems, allowing for a better approximation of the cellular organization and complexity of the developing human brain [30]. Furthermore, the isogenic human-based system ensures a more faithful portrayal of disease processes during neurodevelopment compared with rodent models. We find that ventricular zone-like neuroepithelial progenitor layer expansion is blunted in a *HTT* CAG repeat length-dependent and cell intrinsic manner due to premature neurogenesis in HD cerebral organoids. We further identify impairments in cell cycle regulatory processes, which may collectively contribute to premature neuronal differentiation. Finally, we demonstrate increased activity of the ATM-p53 pathway, an up-stream regulator of cell cycle processes, and show that treatment with ATM antagonists partially rescues the blunted neuroepithelial progenitor expansion in HD organoids.

## Results

### Expanded HTT CAG repeats lead to a reduced number of large neuroepithelial structures in cerebral organoids

To understand the effect of *HTT* CAG repeat length on early neurodevelopment and organization, we generated cerebral organoids from a panel of isogenic HD (IsoHD) hESC lines harboring *HTT* CAG lengths of 30, 45, 65 and 81 repeats, alleles that are associated with control, adult-, adolescent-, and juvenile-onset HD, respectively [1,2]. For each line, we chose 2–3 independent hESC clones with confirmed pluripotency, normal karyotype, and stable maintenance of the *HTT* CAG repeat tracts [31]. The hESC lines were induced to form cerebral organoids using an established protocol [30] whereby suspended hESCs are reaggregated to form embryoid bodies (EB) that grow to form neuroectoderm and contain neuroepithelium that expands over time (Figure 1A). At day 28 of cerebral organoid formation, a time-point which mimics 2–3 months of human in vivo brain development [32], the sizes of the organoids were measured. No difference in organoid size between hESC lines with different CAG lengths was detected (Figure 1B).

**Figure 1.**
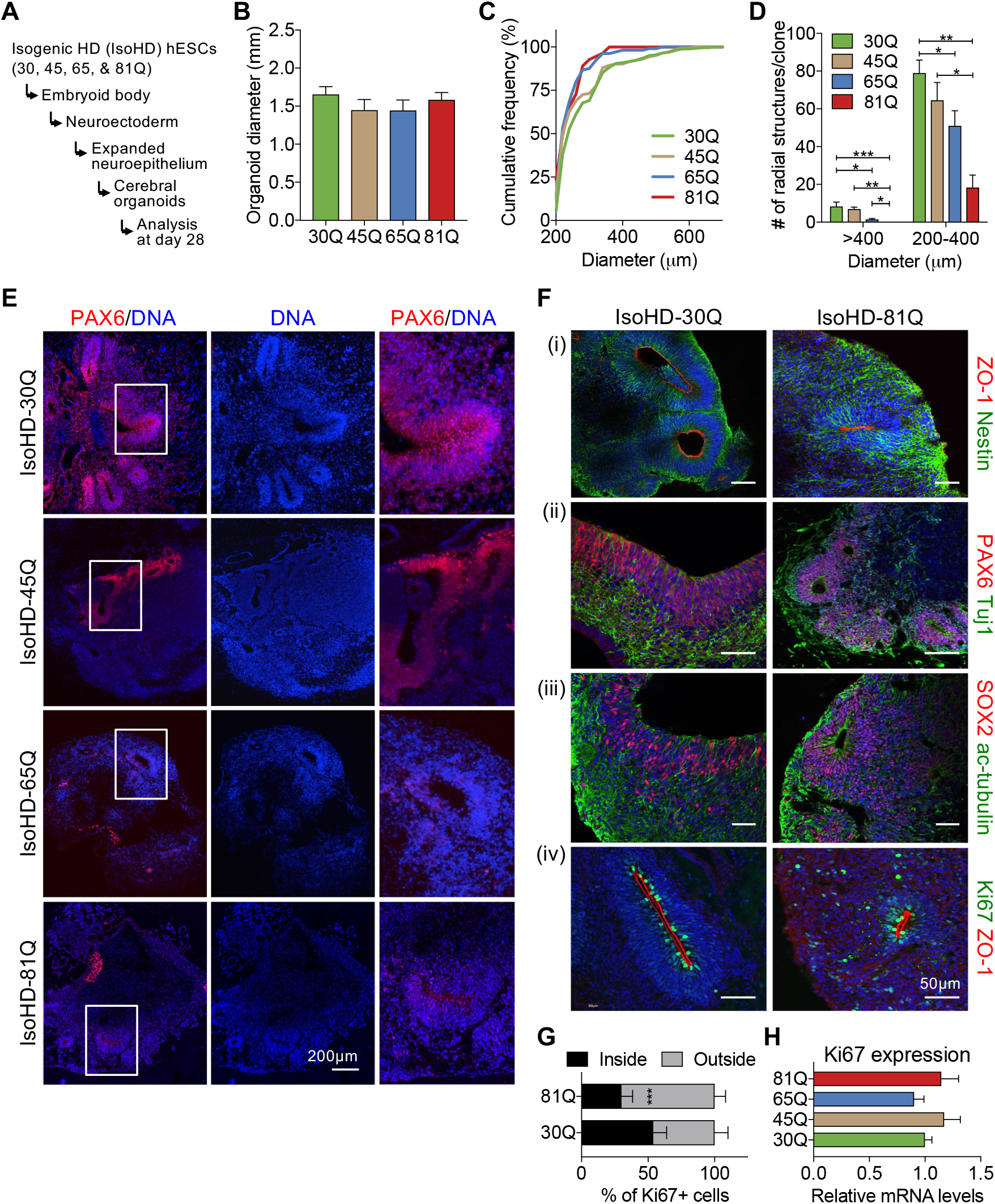
Huntingtin CAG length-dependent impairment in expansion of PAX6-positive neuroepithelial structures in isogenic HD cerebral organoids. **(A)** Analysis of cerebral organoid sizes at day 28 of differentiation. **(B)** For each organoid, the diameters were measured under microscope with three measurements taken at 45 degree angles to obtain the mean value. For each line, there are 2-3 clones, and 6 organoids per clone were counted. Values represent mean ± SEM. **(C)** Organoids were sectioned at 200μm intervals and five continuous sections for each organoid were stained with Hoechst and PAX6. Cumulative frequency analysis of neuroepithelial structures diameters. **(D)** Radially-organized PAX6-positive, neuroepithelial structures greater than 400μm or between 200μm and 400μm were counted. For each line, there are 2-3 clones, and 6 organoids per clone were counted. Values represent mean ± SEM. (*p<0.05, **p<0.01, ***p<0.001; One-way ANOVA with Fisher’s LSD post-hoc test). **(E)** Immunostaining of organoid sections. Representative images of IsoHD-30Q, 45Q, 65Q and 81Q organoids in low magnification showing PAX6-positive radial (neuroepithelial) structures. Insert is enlarged in right panel. Scale bar: 200μm. **(F)** Representative images of organoid staining with neural markers. 81Q organoids showed disorganized staining of PAX6, SOX2, and Ki67. Scale bar: 50μm. **(G)** Percentage of Kİ67+ cells inside and outside radial structures. Values represent mean ± SD. (H) qRT-PCR analysis of Ki67 expression in IsoHD organoids at day 28. (n=3/genotype). Values represent mean ± SD.

A hallmark of early forebrain development is the formation of large continuous neuroepithelial structures with well-defined progenitor zones and neuronal populations [33]. To examine the formation of large radial structures in day 28 organoids, we sectioned each organoid at 200μm intervals and counted the number of radial structures with diameters over 200μm for each IsoHD line. Cumulative frequency analysis showed a shift towards smaller radial structures in IsoHD-81Q and IsoHD-65Q organoids compared with IsoHD-45Q and IsoHD-30Q lines (Figure 1C). In 81Q organoids, radial structures larger than 400 μm were not found, and the number of radial structures with diameters in the 200–400μm range was significantly lower than those found in 30Q control lines (Figure 1D).

The majority of the IsoHD-81Q radial structures were less than 200μm in diameter with immature and rosette-like characteristics. In IsoHD-65Q organoids, the number of large radial structures was also significantly lower than in IsoHD-30Q control organoids, whereas for IsoHD-45Q, large radial structures were frequently seen, similar to controls. The results indicate that the number of radial structures decreases with the expansion of the *HTT* CAG tract in a CAG repeat length-dependent manner.

We then examined day 21 organoids (Figure S1A). At this earlier time-point, organoids with different CAG lengths showed a similar number of radial structures with diameters over 200μm (Figure S1B). Radial structures larger than 400μm were absent in day 21 organoids from all CAG repeat lengths. This indicates that the lack of large radial structures in IsoHD-81Q organoids at day 28 is not due to an inability to form neuroepithelium but rather a failure to expand such structures.

Next, we characterized the cerebral organoids and radial structures using neuroepithelial and neural progenitor markers, focusing on IsoHD-30Q and 81Q organoids. Immunofluorescence staining showed that in day 28 IsoHD-81Q organoids, PAX6+ radial structures were small, immature, and rosette-like (Figure 1E). Progenitor cells in the small IsoHD-81Q radial structures (Figure 1F,i) were lacking in organization with PAX6+ and SOX2+ cells found scattered beyond the radial structures (Figure 1F,ii,iii). Similarly, Ki67-positive proliferating cells appeared to be disorganized and scattered away from the apical surface of the neuroepithelial layer in IsoHD-81Q organoids (Figure 1F,iv, 1G). Despite the altered distribution of Ki67+ cells, qRT-PCR analysis showed comparable levels of Ki67 across all CAG lengths (Figure 1H). The presence of comparable levels of actively proliferating cells in the different IsoHD lines may explain our observation that HD organoids grew to a similar size compared with 30Q control organoids despite the reduced size of radial structures.

With extended culture to 42 days, both 30Q and 81Q organoid express cortical neural marker CTIP2 and the significant difference of radial structures between 30Q and 81Q still remains (Figure S2).

Because specific neural region formation in organoids is highly variable, the experiment was repeated with a short treatment of GSK inhibitor (CHIR99021) and a TGF-beta inhibitor (SB431542) before organoid embedding to promote consistent forebrain tissue formation [34,35]. Immunostaining showed consistent forebrain marker FOXG1 expression in both 30Q and 81Q organoids. qRT-PCR analysis showed expression of dorsal forebrain identity markers *PAX6, FOXG1, OTX2*, and *TBR1* with no expression of ventral identity markers *GSX2* and *NKX2.1* for both IsoHD-30Q and 81Q lines (Figure S3). Consistently, a significant difference in the size of neuroepithelial radial structures between 30Q and 81Q was identified in these forebrain-patterned cerebral organoids (Figure S4).

### HD hiPSC-derived cerebral organoids show abnormal neuroepithelial structures similar to IsoHD lines

To verify the abnormal neuroepithelial formation phenotype observed in IsoHD lines, we next used HD hiPSCs with 18Q, 71Q and 109Q [36] to form cerebral organoids. Control 18Q hiPSC-derived organoids contained large radial structures, some over 600μm in diameter with an expanded layer of PAX6+ and SOX2+ progenitor cells (Figure 2A-C). However, such structures were not found in the 71Q and 109Q HD hiPSC-derived organoids. Most radial structures in the 71Q and 109Q cerebral organoids were smaller than 200 μm in diameter with rosette-like morphology (Figure 2A,C). Similar to IsoHD organoids, a TBR2-positive layer of cells was not found in 71Q and 109Q organoids. However, there were several individual TBR2-positive cells located around small rosette-like structures indicating that the capacity to form TBR2-positive cells in HD organoids was not impaired, and that the absence of a TBR2-positive cell layer was most likely due to the reduced size and abnormal development of radial structures (Figure 2B). *TBR2* mRNA levels were reduced in 109Q cerebral organoids, consistent with the immunostaining findings (Figure 2D). These results indicate that IsoHD hESC- and HD hiPSC-derived cerebral organoids exhibit similar phenotypes, demonstrating impaired formation of large radial structures and neural progenitor organization.

**Figure 2.**
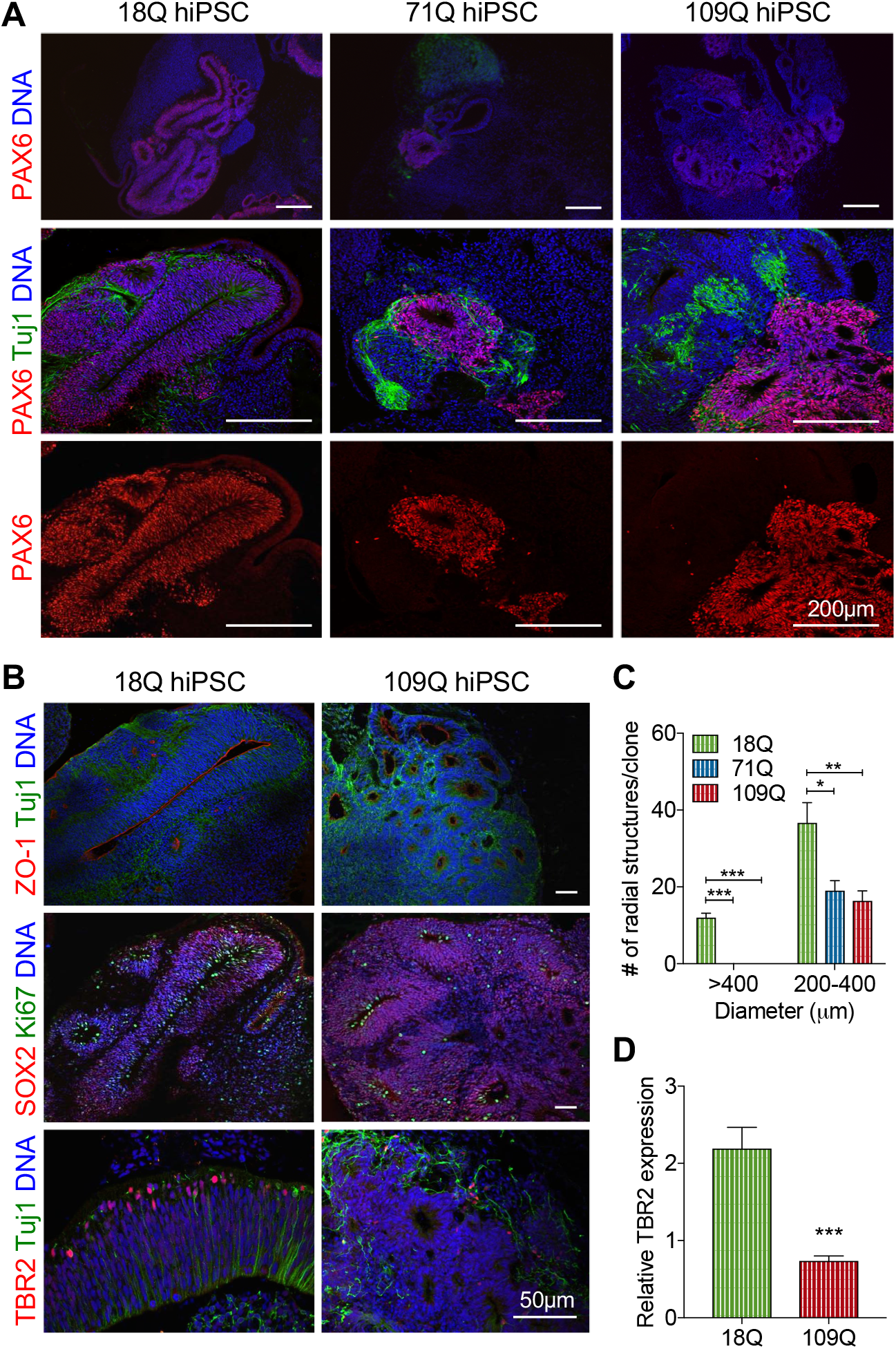
Impaired neural development is seen in HD hiPSC-derived cerebral organoids. **(A)** Immunostaining of hiPSC-derived organoids at day 28 showing an expanded neuroepithelial layer with PAX6-positive cells in 18Q-hiPSCs, but disorganized PAX6 staining in 71Q and 109Q hiPSCs. **(B)** Representative images of organoid staining with neural markers. (C) Organoids were sectioned at 200μm intervals and five continuous sections for each organoid were stained with Hoechst and PAX6. Radially organized neuroepithelial structures greater than 400μm or between 200μm and 400μm were counted. For each line, six organoids per line were counted. 71Q and 109Q organoids were devoid of 400-μm structures and had significantly a lower number of 200-400μm structures than 18Q control organoids. Values represents mean ± SEM. **(D)** Relative *TBR2* mRNA levels. Values represents mean ± SEM. *p<0.05, **p<0.01, ***p<0.001, One-way ANOVA with Fisher’s LSD post-hoc test.

### Cell-intrinsic mechanisms drive the abnormal development of neuroepithelial structures in IsoHD-81Q cerebral organoids

To investigate whether abnormal neuroepithelial formation in IsoHD-81Q cerebral organoids reflects cell-intrinsic mechanisms, we performed a chimerism experiment in which IsoHD-81Q hESCs labeled with GFP were co-cultured in a 1:1 ratio with unlabeled IsoHD-30Q hESCs and then induced to form forebrain organoids (Figure 3A). At day 28, large radial structures were formed by the co-cultured cells. However, close examination revealed that large radial structures over 400μm in diameter were formed solely of IsoHD-30Q cells and were devoid of IsoHD-81Q-GFP+ cells (Figure 3B,i). This observation suggests an intrinsic inability of IsoHD-81Q cells to form large radial structures. In chimeric radial structures containing 81Q-GFP+ cells, we observed a disturbed and disorganized neuroepithelial layer with many PAX6+ cells located away from the VZ-like layer (Figure 3B,ii). The disorganization was more pronounced in structures with more integrated 81Q-GFP+ cells (Figure 3B,iii). The decrease of PAX6 positive progenitor cells in radial structures was highly correlated with the increase of GFP positive cells (Figure 3C,i). To rule out the contribution of GFP-labeling to the IsoHD-81Q phenotype, we labeled IsoHD-30Q cells with GFP and formed chimeric organoids with unlabeled IsoHD-30Q cells. The formed chimeric organoids confirmed that GFP-labeled IsoHD-30Q cells maintained their ability to form large neuroepithelial structures (Figure S5). Furthermore, there was no significant correlation between PAX6 positive and GFP positive progenitor cells in radial structures (Figure 3C,ii). Taken together, these findings suggest that the blunted radial structures in IsoHD-81Q cerebral organoids result from cell autonomous deficits in the ability of IsoHD-81Q cells to form expanded neuroepithelia.

**Figure 3.**
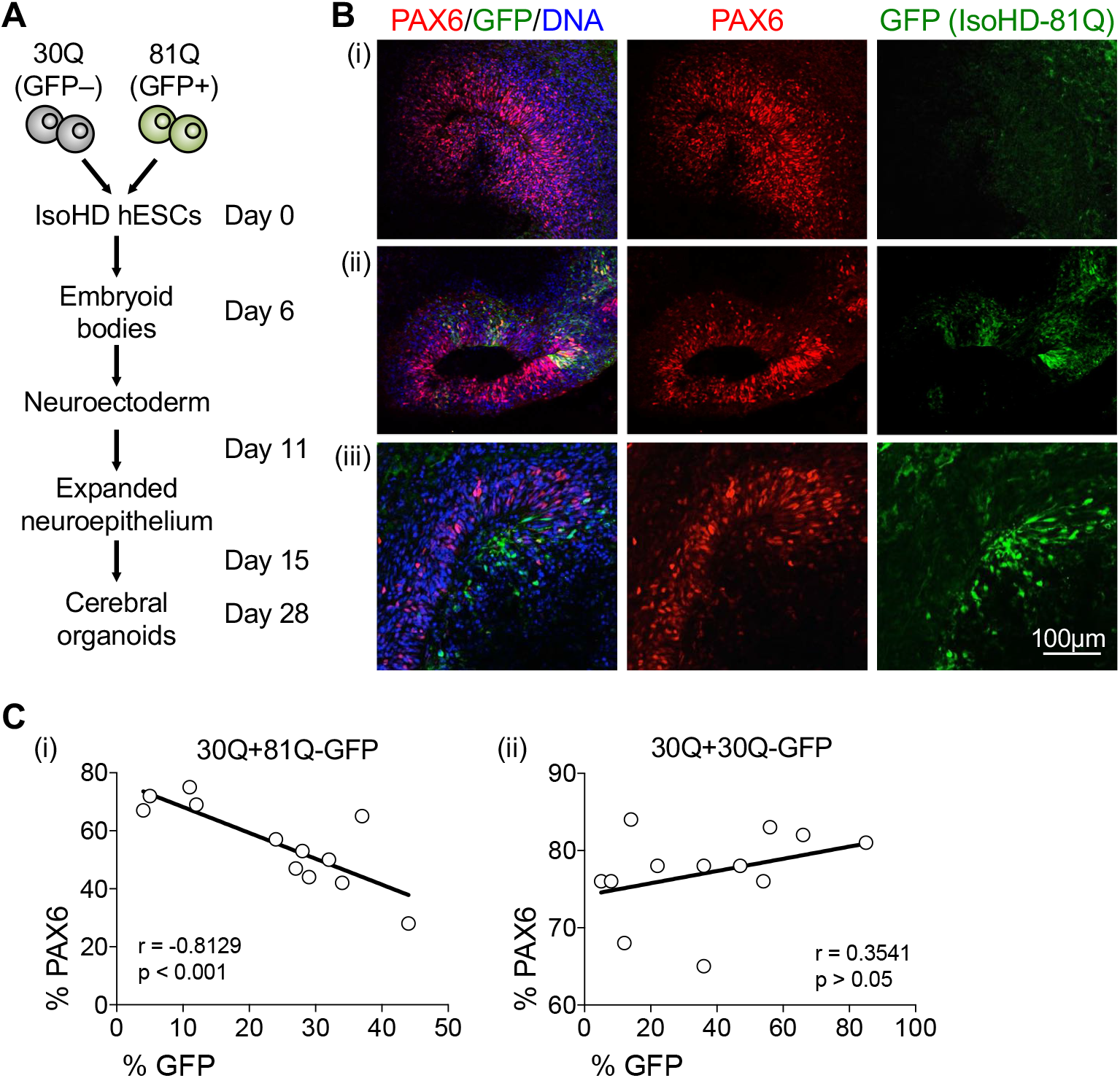
Cell intrinsic mechanisms drive the abnormal development of neuroepithelial structure in IsoHD-81Q cerebral organoids. **(A)** Schematic of IsoHD-30Q/81Q chimerism experiment. GFP-labeled IsoHD-81Q cells were mixed in equal numbers with control vector-infected IsoHD-30Q cells, and organoids were analyzed at day 28. **(B)** Neuroepithelial development in chimeric IsoHD-30Q/81Q organoids. Organoid sections were stained for GFP and PAX6. **(C)** Correlation of GFP-positive cells with PAX6-positive cells in chimeric radial structures from (i) 81Q-GFP co-culture with 30Q and (ii) 30Q-GFP co-culture with 30Q. 12 chimeric radial structures were examined for co-cultured organoids. The percentage of GFP-positive and PAX6-positive cells were calculated for each radial structure. Pearson correlation coefficient (r) analysis shows significant correlation for 30Q+81Q-GFP (i) but not 30Q-GFP+30Q (ii) chimeric organoids.

### RNA-seq analysis reveals disruption of key cell cycle regulatory pathways in IsoHD-81Q cerebral organoids

To understand the cellular mechanisms underlying the abnormal neurodevelopment in IsoHD organoids, we performed RNA-seq analysis on IsoHD-30Q and 81Q organoids at day 28 of differentiation (Figure 4). Three separate clonal lines each were used for IsoHD-30Q and 81Q, and three organoids were pooled for each clonal line. Multi-dimensional scaling (MDS) of the gene expression data revealed a clear separation in transcriptional profiles between the IsoHD-30Q control and the 81Q HD organoids (Figure 4A). We identified 1,890 differentially expressed genes (DEGs; false discovery rate (FDR) <10%) between IsoHD-30Q and 81Q, of which 773 genes were downregulated and 1,117 genes were upregulated in 81Q organoids, as depicted in the volcano plot (Figure 4B).

**Figure 4.**
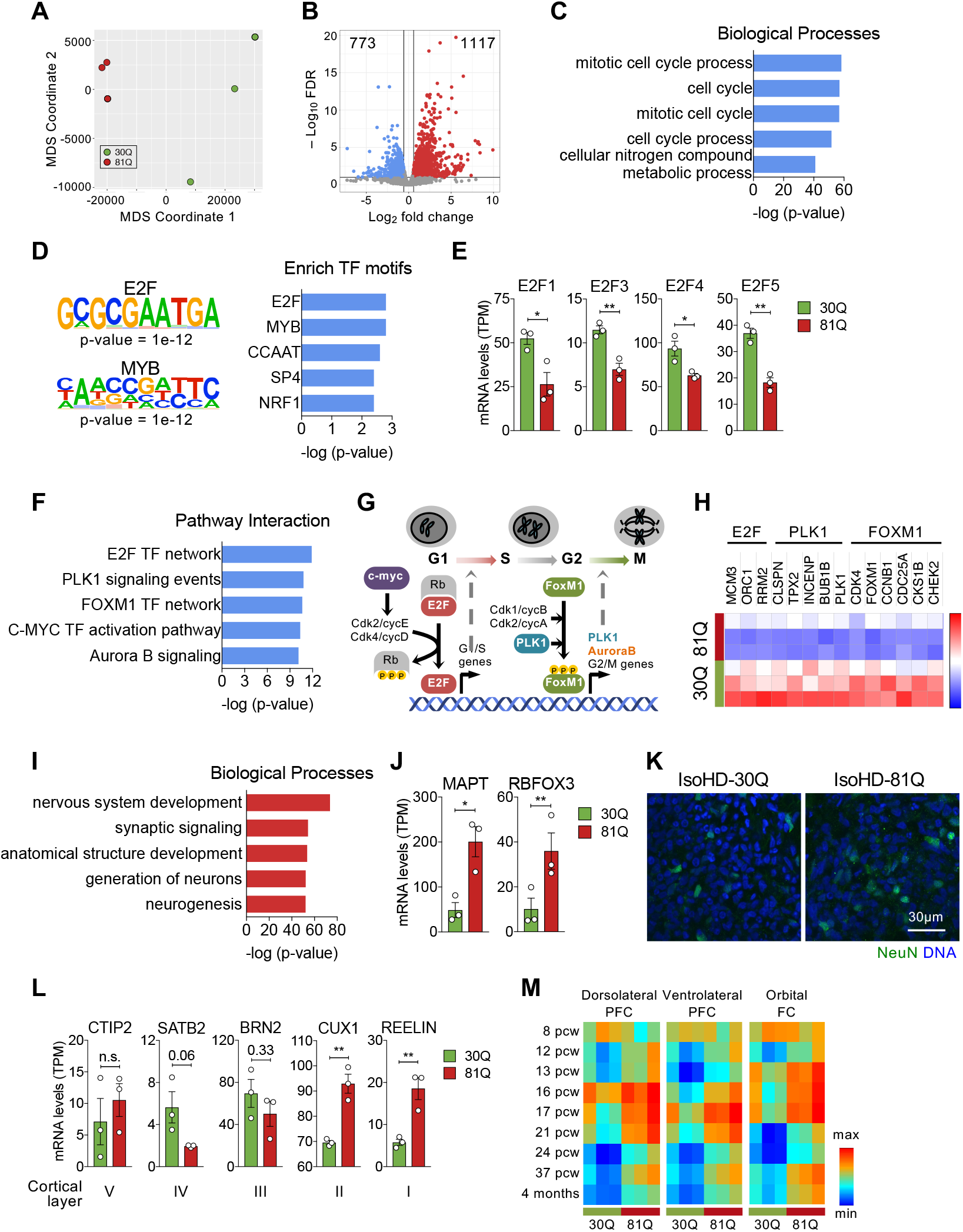
RNA-seq analysis of organoids reveals disruption of key cell cycle pathways and premature neurogenesis in 81Q cerebral organoids. **(A)** MDS plot showing separation of the IsoHD-30Q and IsoHD-81Q samples. Three separate lines were used for 30Q and 81Q, and three organoids were pooled per line. **(B)** Volcano plot showing the differentially expressed genes between IsoHD-81Q and IsoHD-30 cerebral organoids. **(C)** Gene ontology analysis of down-regulated genes in IsoHD-81Q organoids. The top five significant terms for downregulated genes are shown. **(D)** E2F and MYB appear as the top motifs enriched in DEGs down-regulated in IsoHD-81 compared with IsoHD-30Q organoids. **(E)** Levels of several members of the E2F TF family in IsoHD-30Q and IsoHD-81 Q organoids. Two-tailed t-test; *p<0.05; **p<0.01; n=3/group. **(F)** Annotation of down-regulated genes using the Pathway Interaction Database suggests disruption of key cell cycle regulatory pathways in IsoHD-81Q organoids. **(G)** Diagram depicting cell cycle regulatory pathways disrupted in IsoHD-81Q organoids. **(H)** A heatmap of levels of representative genes from the E2F, PLK1, and FOXM1 pathways. (I) Functional annotation of up-regulated genes shows enrichment of neurogenesis and neuronal maturation-related processes in 81Q organoids. **(K-L)** Consistent with enrichment analysis, levels of neuronal maturation **(J,K)** and cortical neuron development markers **(L)** are increased in 81Q organoids. Two-tailed t-test; *p<0.05; **p<0.01. n=3/group. **(M)** Heatmaps of Pearson’s correlation analysis comparing genes expressed in IsoHD-30Q and IsoHD-81Q with published gene expression profiles of human fetal brain (8 pcw-4 mo). IsoHD-81Q cerebral organoid gene expression profile shows higher similarity with late stage brain development (16-21 pew) compared with 30Q cerebral organoids.

Functional annotation of genes downregulated in IsoHD-81Q organoids using gene ontology (GO) analysis showed an enrichment for GO terms related to mitosis and cell cycle processes (Figure 4C). To examine whether certain DNA binding site motifs were enriched in the downregulated DEGs, we applied a motif-discovery algorithm, HOMER [37]. Consistent with the enrichment of DEGs for cell cycle genes, E2F and MYB consensus-binding motifs were identified as the top two motifs enriched amongst genes downregulated in IsoHD-81Q organoids (Figure 4D). The transcription factors (TF) E2F and MYB play a key role in cell cycle control [38,39]. Expression levels of several members of the E2F TF family such as *E2F1, E2F3, E2F4* and *E2F5* were significantly lower in IsoHD-81Q compared with 30Q organoids (Figure 4E). Further annotation using the Pathway Interaction Database [40], a curated collection of regulatory and signaling pathways, corroborated this finding (Figure 4F). The top enriched pathways amongst down-regulated genes included E2F, FOXM1, and c-Myc TF networks, along with PLK1 and Aurora B signaling, all pathways that play key roles in different phases of the cell cycle (Figure 4G). A heatmap gene expression from representative genes in these pathways is shown in Figure [40].

### Premature neurogenesis in IsoHD-81Q cerebral organoids

We next examined the profile of genes up-regulated in IsoHD-81Q compared with IsoHD-30Q cerebral organoids. Surprisingly, GO analysis showed enrichment of neurogenesis and neuronal development-related processes (Figure 4I). Consistent with the enrichment analysis, levels of the neuronal maturation markers MAPT (encoding tau) and RBFOX3 (encoding NeuN) were significantly increased in IsoHD-81Q cerebral organoids (Figure 4J). NeuN immunostaining in IsoHD-81Q organoids showed a higher percentage of positive cells than that of IsoHD-30Q (Figure 4K). In addition, expression levels of cortical layer I and II markers, *CUX1* and *REELIN*, were significantly higher in IsoHD-81Q organoids compared with IsoHD-30Q (Figure 4L). Furthermore, we compared the transcriptome of IsoHD-30Q and 81Q organoids with published gene expression profiles of human fetal brain from three different cortical regions ranging from 8 pcw (post-conception week) to 4 months from the Allen Brain Atlas. The results showed that the transcriptome of IsoHD-81Q cerebral organoids was more closely correlated with later stages of human in vivo brain development (16–21 pcw) than that of IsoHD-30Q (Figure 4M). Collectively, these results point to premature neurogenesis in the IsoHD-81Q as compared with the IsoHD-30Q cerebral organoids.

### Increased G1 length in IsoHD-81Q neural progenitors

The importance of cell cycle regulation in neural development is well established [41,42]. As neural progenitors become neurogenic, their cell cycle lengthens due to an extended G1 phase [43]. These changes in the cell cycle are key regulators of fate specification during early neurogenesis. Indeed, increasing the length of the cell cycle by down-regulating cdk activity promotes neuronal differentiation [44]. Conversely, shortening of the cell cycle by overexpression of cdk4/cyclinD1 promotes neural progenitor proliferation and expansion and delays neurogenesis [45].

We reasoned that premature neurogenesis in IsoHD-81Q cerebral organoids may be partly due to increased G1 length. To test this hypothesis, we used the Fluorescent Ubiquitination-based Cell Cycle Indicator (FUCCI) assay [46] to label neural stem cells derived from IsoHD-30Q and 81Q in different phases of the cell cycle (Figure 5A). Time-lapse recording of the cell cycle revealed that IsoHD-81Q neural stem cells exhibited a significantly longer G1 duration than 30Q (Figure 5B). No difference in the length of the S/G2-M phases was observed between IsoHD-81Q and 30Q neural stem cells (Figure 5C).

**Figure 5.**
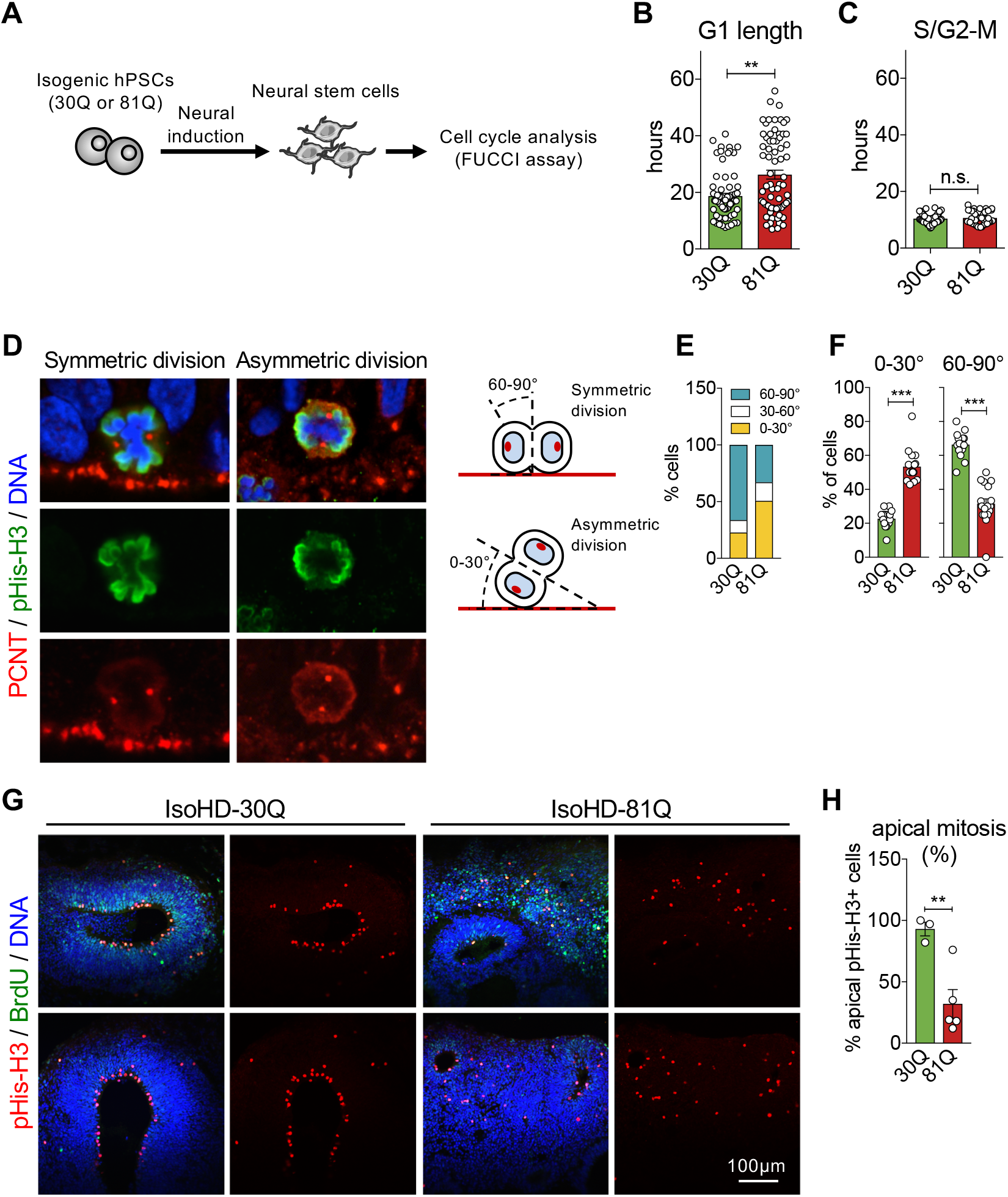
Increased G1 length and reduced symmetric apical division of neural progenitors in IsoHD-81Q cerebral organoids. **(A-C)** The cell cycle was analyzed in neural stem cells derived from IsoHD-30Q and 81Q hiPSCs using the FUCCI assay **(A).** The length of the G1 phase was found to be increased in IsoHD-80Q neural stem cells compared with IsoHD-30Q **(B).** No difference in S/G2-M was found between IsoHD-30Q and 80Q **(C).** Mann-Whitney U test, **p<0.01. n.s.= not significant. **(D-F)** Analysis of symmetric versus asymmetric division in IsoHD-30Q and 81Q organoids at day 28. **(D)** Immunostaining of pericentrin (PCNT, red) and phospho-histone H3 (Serine 28) (pHis-H3, green) showing examples of symmetric and asymmetric apical division observed in cerebral organoids. Only apical mitotic cells were considered for analysis of the cleavage angle. **(E)** Quantification of symmetric and asymmetric divisions. Collectively 72-85 dividing cells were analyzed from IsoHD-30Q and IsoHD-81Q organoids. **(F)** Significantly more asymmetrically dividing apical cells were identified in IsoHD-81Q radial structures, whereas more symmetrically dividing cells were identified in IsoHD-30Q radial structures. Mann-Whitney U test, ***p<0.001. **(G,H)** Immunostaining of IsoHD-30Q and 81Q organoids for the mitosis marker pHis-H3 (red) and Brdll (green) at day 28 **(G)** shows significantly less apical mitosis in IsoHD-81Q compared with IsoHD-30Q cerebral organoids **(H).** Two-tailed t-test; **p<0.01.

### Reduced symmetric division of apical progenitors in IsoHD-81Q cerebral organoids

During embryonic corticogenesis, a single layer of neuroepithelial progenitors is formed that subsequently expands through symmetric divisions [33]. Following an initial expansion phase, the division of neuroepithelial progenitors becomes more asymmetric, leading to the generation of intermediate progenitors that move away from the ventricular zone. Intermediate progenitors subsequently give rise to post-mitotic neurons through further asymmetric division [47].

We hypothesized that the cell cycle impairment in IsoHD-81Q organoids may skew apical cell division of neuroepithelial stem cells to be more asymmetric, precipitating smaller radial structures in IsoHD-81Q cerebral organoids. We thus examined the proportion of apical progenitors undergoing symmetrical and asymmetrical division by dual staining for the mitotic cell marker phospho-histone 3 (serine-28; pHis-H3) and pericentrin (PCNT), a component of the centrosomes which also marks the apical neuroepithelial surface. Progenitor cells undergoing symmetrical division were identified by a division angle of 60–90° whereas those undergoing asymmetric division were identified by a division angle of 0–30° (Figure 5D). Assessment of division angle identified a significantly higher percentage of symmetrical apical division, which is associated with neuroepithelial expansion in IsoHD-30Q control organoids (Figure 5E,F). Conversely, we observed a higher percentage of asymmetrical apical division, which is associated with a neurogenic switch, in the IsoHD-81Q organoids (Figure 5E,F). Consistent with the increase in asymmetric apical division, we observed a reduced percentage of proliferating apical (pHis-H3+) cells in IsoHD-81Q organoids compared with IsoHD-30Q (Figure 5G,H). These findings indicate that reduced symmetric division may contribute to the blunted expansion of neuroepithelium in HD cerebral organoids.

### Elevated activity of the ATM-p53 pathway contributes to the impairment in neuroepithelium expansion in HD cerebral organoids

The protein kinase ataxia telangiectasia mutated (ATM), a member of the phosphoinositide 3-kinase (PI3K)-related protein kinase family, is a key up-stream regulator of cell cycle processes [48]. ATM is activated in response to DNA damage, oxidative stress, and altered chromatin structure, and phosphorylates multiple targets, ultimately resulting in the activation of the p53 pathway and downstream cell cycle control processes. A recent study showed that ATM activity is elevated in HD animal and cell models [49]. Thus, we reasoned that the dysregulation of cell cycle processes in IsoHD-81Q cerebral organoids may be, in part, due to elevated ATM-p53 signaling. To test this, we examined the ATM-p53 pathway by immunoblotting control and HD cerebral organoids. We observed an increase in the levels of pKAP1 (Serine-824), a specific marker of ATM-dependent pathway activation, in IsoHD-81Q organoids compared with IsoHD-30Q (Figure 6A). We further observed stabilization of p53 protein levels in IsoHD-81Q cerebral organoids that correlated with upregulation of p21 protein expression, indicating an activation of the p53/p21 pathway (Figure 6A). Together, these findings indicate that aberrant activation of the ATM-p53 pathway may contribute to cell cycle impairment, including lengthening of the G1 phase, blunted neuroepithelial expansion, and premature neurogenesis in IsoHD-81Q cerebral organoids.

**Figure 6.**
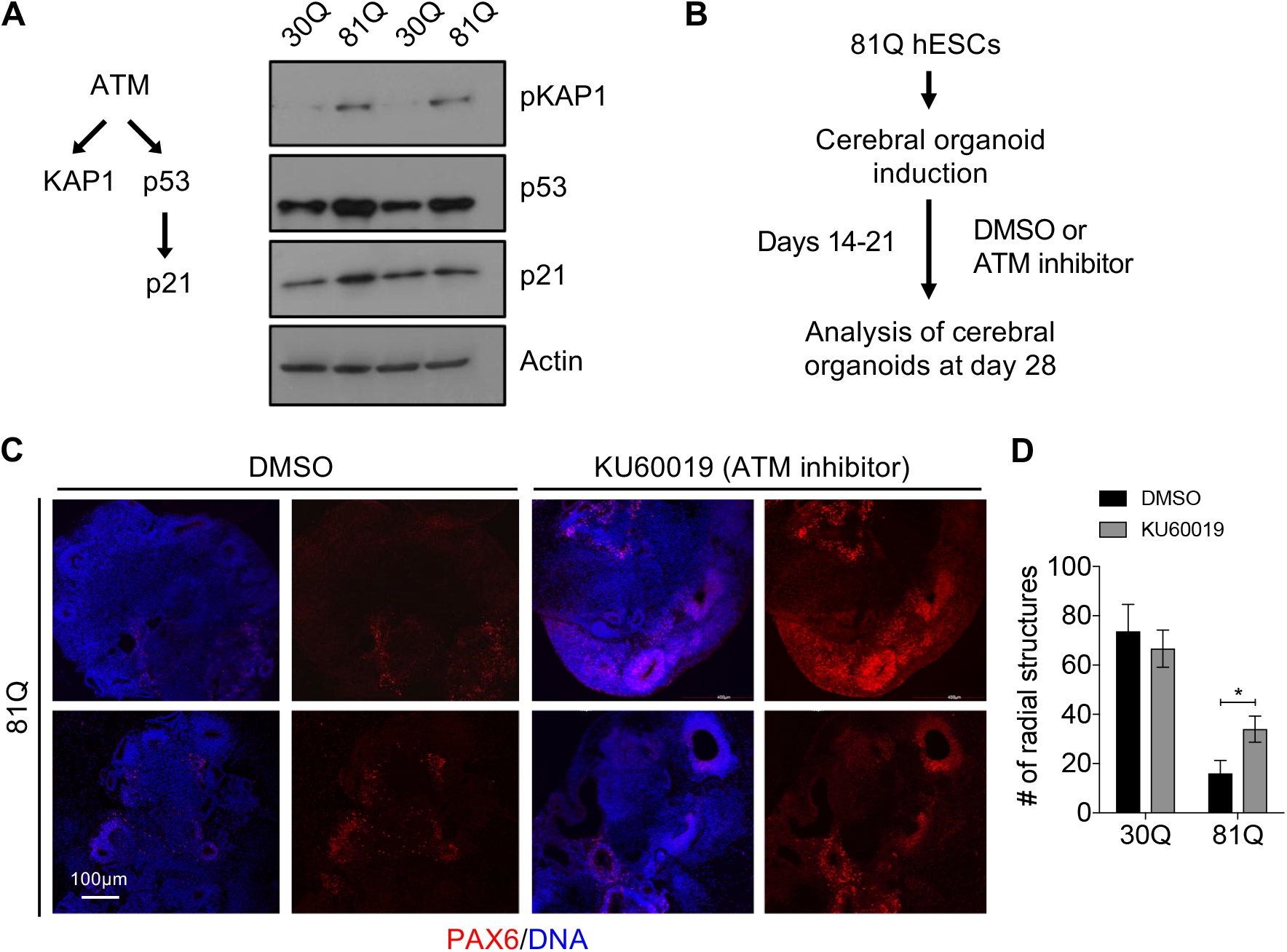
Elevated activity of the ATM-p53 axis contributes to the impairment in neuroepithelium expansion in HD cerebral organoids. **(A)** Western blot analysis of pKAP1, p53, and p21 levels, markers of ATM-dependent pathway activation in IsoHD-30Q control and IsoHD-81Q cerebral organoids. **(B)** Schematic showing induction and differentiation of IsoHD-81Q cells into HD cerebral organoids using the ATM inhibitor KU60019. **(C)** Immunostaining of KU60019-treated IsoHD-81Q organoids showing improved neuroepithelial progenitor layer expansion. **(D)** Quantification of neuroepithelial layers showed a significantly higher number of radial structures >200μm in diameter in IsoHD-81Q organoids treated with KU60019 compared with DMSO. There was no effect of KU60019 treatment on the number of radial structures in IsoHD-30Q organoids. Two-tailed t-test; *p<0.05.

To test whether moderation of ATM activation can rescue the impairment in neuroepithelium layer formation in HD cerebral organoids, we treated IsoHD-81Q cells with KU60019, an ATM inhibitor [50], from days 14 to 21 of differentiation and analyzed the cerebral organoids at day 28 (Figure 6B). We observed significantly improved neuroepithelial progenitor layer expansion in KU60019-treated cerebral organoids compared with DMSO-treated organoids (Figure 6C,D). These results support a role for ATM pathway hyperactivation in the abnormal neurodevelopment of HD cerebral organoids.

## Discussion

Investigating the impact of human genetic variation on neurodevelopmental processes, in particular corticogenesis, using model systems has been challenging to date. This is in part due to the stark differences in the developmental demands between the gyrencephalic human brain and lissencephalic rodents commonly used for disease modeling [51]. Here, we used human cerebral organoids which allow better approximation of the complexity of the developing human brain to investigate the impact of mutant HTT on early neurodevelopment. We show that ventricular zone-like neuroepithelial progenitor layer expansion is blunted in an *HTT* CAG repeat length-dependent manner. We further demonstrate that mutant HTT impairs cell cycle regulatory processes, leading to increased G1 length along with asymmetric division of neuroepithelial progenitors, resulting in premature neuronal differentiation. Finally, we demonstrate increased activity of the ATM-p53 pathway, an up-stream regulator of cell cycle processes [48], and show that treatment with ATM antagonists partially rescues the blunted neuroepithelial progenitor expansion in HD organoids. Overall, our findings suggest that *HTT* CAG repeat length regulates the balance between neural progenitor expansion and differentiation during early neurodevelopment (Figure 7).

**Figure 7.**
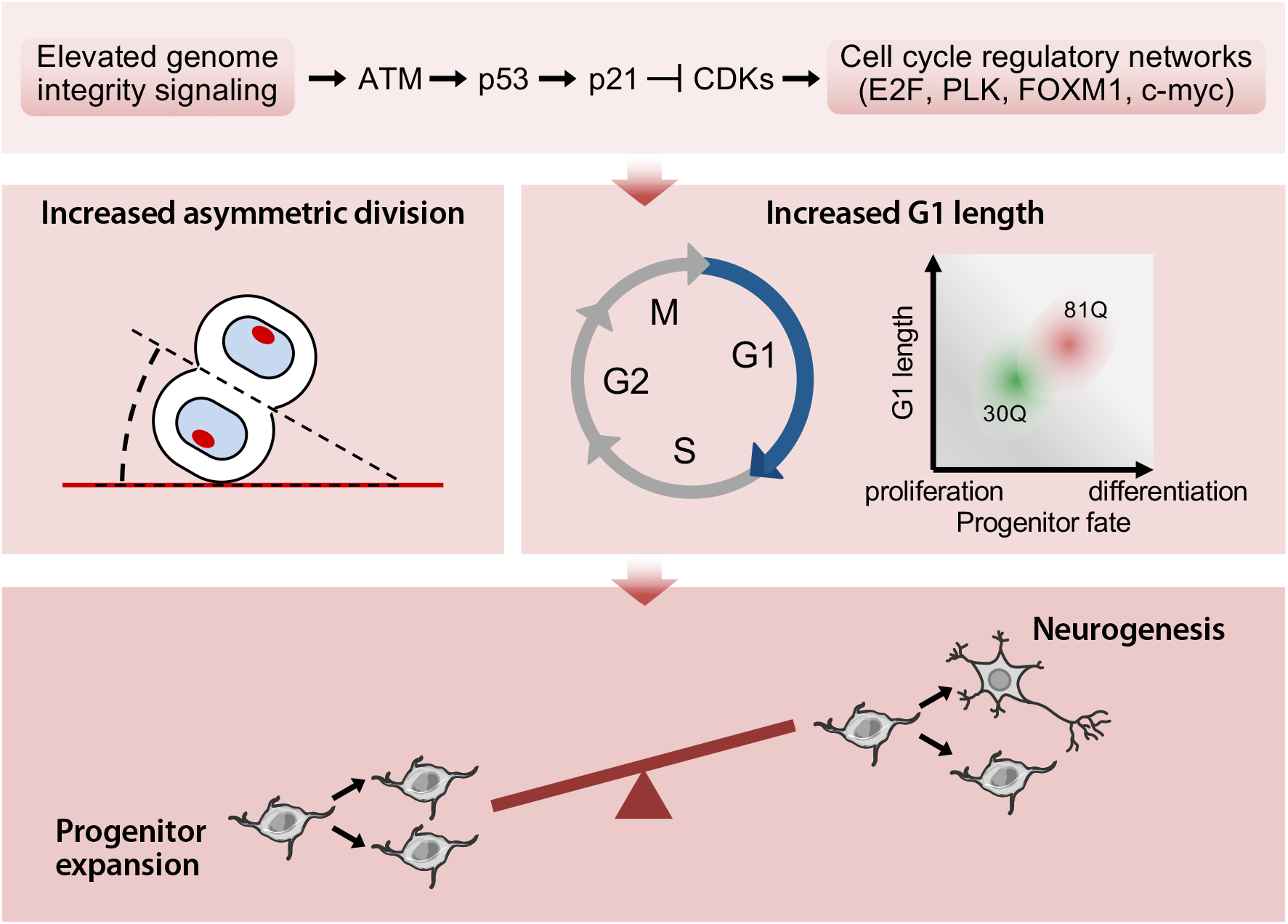
Model of mechanisms contributing to premature neurogenesis in HD cerebral organoids. Elevated genome integrity signaling in HD cerebral organoids, likely triggered by a combination of DNA damage, oxidative stress, and altered chromatin structure due to the large expansions of *HTT*’s CAG trinucleotide repeats, impairs cell cycle regulatory processes, particularly the G1-to-S promoting E2F and c-myc-related pathways, leading to increased G1 length. Concomitantly, altered cleavage angle during mitosis leads to increased asymmetric division of apical progenitors in HD cerebral organoids. Combined, the increased G1 length and asymmetric apical division result in reduced neuroepithelial progenitor expansion and premature neurogenesis in HD organoids. Overall, these ndings suggest that CAG repeat length regulates the balance between neural progenitor expansion and differentiation during early neurodevelopment.

Previous studies examining the role of normal HTT as a scaffold and microtubule-binding protein have shown that it localizes to the poles of mitotic spindles and regulates their orientation during cell division [12]. In line with and supporting our findings, mutant HTT was found to cause spindle misorientation and alter cleavage angle and neural progenitor fate specification in mice [13, 20]. The mechanisms proposed to underlie these abnormalities include altered recruitment and attachment of the motor proteins involved in the alignment and movement of spindle poles during cell division [20] as well as elevated Notch signaling [13], a pathway known to promote asymmetric division of neural progenitors [52]. Here we present evidence that elevated ATM-p53 signaling contributes to the altered neural progenitor proliferation/differentiation balance in HD. ATM signaling plays an important role in neural cell cycle control and neurogenesis [53]. ATM was shown to regulate key cell cycle checkpoints and loss of its activity results in increased proliferation of neural progenitors and impaired neurogenesis in ATM KO mice [53]. Our results using hESC-derived cerebral organoids demonstrate that under conditions of elevated ATM activity, a converse phenomenon is observed resulting in blunted expansion of neural progenitors and accelerated neurogenesis.

Elevated ATM signaling has been documented in models of HD [49,54,55], although the triggers of this increased activity remain unclear. ATM is activated in response to oxidative stress and DNA damage [48], and there is evidence supporting a role for these stressors in HD. Expression of mutant HTT has been shown to result in elevated levels of reactive oxygen species [55], in part due to reduced expression of antioxidant enzymes [54], which may contribute to ATM activation. Similarly, accumulation of single and double-strand DNA breaks in response to mutant HTT expression has been shown to cause increased ATM activation [56]. Consistent with these observations, we observe CAG repeat length-dependent increases in reactive oxygen species as well as activation of ATM in response to DNA damage in neural progenitors derived from the IsoHD hESC panel [31].

In addition to DNA damage and oxidative stress, altered chromatin structure can also lead to partial ATM activation [48]. A recent study has shown that repeat expansion disease genes, including HTT, localize to boundaries of chromatin domains associated with defined chromatin topology [57], The study provided evidence in Fragile X syndrome, a disease caused by CGG expansions in the *FMR1* gene, that pathological repeat expansions are associated with disrupted chromatin boundaries and, by extension, genomic topography [57]. Thus, it is conceivable that the *HTT* CAG repeat expansion in HD also leads to altered chromatin structure, triggering partial ATM activation and elevated genome integrity signaling.

Whether these mutant HTT-mediated alterations in neurodevelopment contribute to the clinical manifestations of the disease is unclear. A recent study using a conditional mouse model of HD has attempted to address this question [58]. Expression of mutant HTT was restricted to development and terminated at postnatal day 21. This selective exposure to mutant HTT during development was sufficient to precipitate a number of HD-related phenotypes, including enhanced susceptibility to excitotoxicity, striatal degeneration, and progressive motor coordination deficits [58]. These findings suggest that developmental effects of mutant HTT may contribute to the pathogenesis of HD, and highlight the importance of a detailed understanding of the role of HTT during development as well as perturbations to its function when mutated. Notably, the *HTT* CAG repeat expansions in the range associated with juvenile onset (>60 repeats) appear to elicit neurodevelopmental abnormalities most robustly both in our study as well as the aforementioned mouse studies. Overall, these findings suggest that HD, at least in its early-onset juvenile forms, may not be a purely neurodegenerative disorder and that abnormal neurodevelopment may be a component of its pathophysiology. This notion is further supported by several studies in children and adolescent HD gene carriers who are estimated to be decades from onset, showing altered intracranial volume [4] and brain region morphology [5,6].

Our findings of premature neurogenesis and neuronal differentiation in HD organoids contrast with those of a recently published study examining the impact of mutant HTT on neurodevelopment using HD hiPSCs [29]. The study reports that HD cortical organoids exhibit a gene expression signature indicative of less mature neuronal development compared to control organoids, concluding that mutant HTT precludes normal neuronal fate acquisition. Despite the differences in findings and conclusions, some similarities exist between the two studies such as the observation of reduced PAX6-positive neuroepithelial cells in HD organoids.

Given that *HTT* CAG repeat expansions can have severe pathological and fatal consequences, it is puzzling that CAG repeat expansions have not only been retained but favored over evolutionary times. This paradox has been suggested to reflect a beneficial effect of CAG repeat expansions on, amongst other things, nervous system development [59]. This notion is largely based on the observation that organisms with more developed nervous systems harbor *HTT* alleles with longer CAG repeats [60]. Indeed, humans have the longest known *HTT* CAG repeat lengths. Our findings now provide a compelling link between neuroepithelium expansion, a cardinal aspect of corticogenesis, and *HTT* CAG repeat length, highlighting an aspect of HTT biology of potential evolutionary significance.

## Materials and Methods

### Human ESC and iPSC lines

The IsoHD female hESC lines and HD hiPSC lines (18Q, 71Q, and 109Q) were previously reported [31,36,61]. The lines were grown and expanded in feeder-free conditions using mTeSR1 medium (STEMCELL Technologies), at 37°C with 5% CO2. Medium was replaced daily.

### Maintenance of IsoHD hESC lines

Human IsoHD lines 30Q, 45Q, 65Q, and 81Q were generated in-house and were fully characterized [31]. Cell lines were maintained with mTeSR medium. Culture medium was changed every day and cells were passaged every week onto a new plate pre-coated with Matrigel. IsoHD lines were detached using 1 mg/ml Dispase (Invitrogen), and were further dissociated by manual pipetting.

### Generation of forebrain organoids

To generate forebrain-specific organoids, a modification of the protocol by Lancaster and colleagues was applied [30]. Human IsoHD cells were detached using Accutase and plated in 96-well ultra-low attachment plates (Corning Costar) at 9×10^3^ cells/well in DMEM/F12 with 20% KnockOut Serum Replacement, ROCK inhibitor (10μM), and low-bFGF (4 ng/ml). On day 5–6, half of the culture volume was exchanged into induction medium (DMEM:F12 containing 1× N2 supplement (Invitrogen), 1× non-essential amino acids, 1× GlutaMax, 10 μg/ml heparin (Sigma), and 1× penicillin/streptomycin). On day 11, organoids were quality-checked. Organoids showing clearing of embryoid body borders and formation of radially organized neuroepithelium were embedded in Matrigel (BD Biosciences). Embedded organoids were transferred to a 6-well low attachment plate and maintained in organoid differentiation medium (DMEM:F12 containing 1× N2 and B27 supplements (with no vitamin A, Invitrogen), 1× penicillin/streptomycin, 1× 2-mercaptoethanol, 1× non-essential amino acids, and 2.5 μg/ml insulin (Sigma)). To promote consistent forebrain tissue formation, organoids were treated with 1 μM GSK inhibitor (CHIR99021) for three days before embedding in Matrigel. On day 14, plates containing organoids were incubated with shaking (70 rpm) in differentiation medium containing vitamin A and B27 supplement. Media were changed every other day.

### Tissue preparation and immunohistochemistry

For fixation, organoids were incubated in 4% paraformaldehyde (PFA) for 30–60 min at room temperature, followed by three washes with phosphate-buffered saline (PBS). The organoids were then incubated overnight in a 30% sucrose solution. Organoids were embedded in gelatin/sucrose solution, frozen in cold isopentane, and sectioned with a cryostat (Leica). For immunostaining, slides were permeabilized with 0.2% Triton-X and blocked in 4% goat/donkey serum in PBS for 1 h. Primary antibodies were diluted in blocking solution and applied to the sections overnight. After washing with PBS-Tween (PBST), secondary antibodies were diluted in blocking solution and applied to the sections for 1 h at room temperature. Finally, sections were washed with PBST and stained with DAPI. All images were acquired an on Olympus FV1000 inverted confocal microscope. The antibodies and dilutions used are listed in Table 1.

**Table 1.** List of antibodies and dilutions used for immunocytochemistry and immunoblotting analysis.

| Antibody | Species | Vendor | Cat. No. | Dilution |
| --- | --- | --- | --- | --- |
| CTIP2 | Rt | Abcam | ab18465 | 1:300 |
| FOXG1 | Rb | Abcam | ab18259 | 1:1000 |
| GFP | Ms | Santa Cruz | sc9996 | 1:300 |
| Ki67 | Ms | Millipore | mab4190 | 1:500 |
| Nestin | Ms | Millipore | mab5326 | 1:500 |
| NKX2.1 | Rb | Santa Cruz | sc-13040 | 1:300 |
| PAX6 | Rb | Covance | prb-278p | 1:500 |
| SATB2 | Ms | Abcam | ab92446 | 1:300 |
| SOX2 | Rb | Abcam | RB5603 | 1:500 |
| SOX2 | Gt | Santa Cruz | sc-17320 | 1:100 |
| TBR2 | Rb | Abcam | ab23345 | 1:300 |
| Tuj1 | Ms | Covance | MMS-435P | 1:1000 |
| Zo-1 | Rb | Abcam | Ab59720 | 1:500 |
| Acetyl-tubulin | Ms | Sigma | T7451 | 1:500 |
| p53 | Ms | Santa Cruz | sc-126 | 1:1000 |
| p21 | Ms | Cell Signaling | 2946 | 1:1000 |
| pKAP-1 | Rb | Abcam | ab70369 | 1:500 |

### RNA isolation and quantitative PCR

RNA was extracted from cells using an RNeasy mini kit (Qiagen) according to the manufacturer’s instructions. cDNA was generated using the Superscript^®^ II Reverse Transcription Kit (Life Technologies) (20μl of cDNA per 1 μg of RNA). To perform quantitative PCR (qPCR), cDNA was diluted ten-fold and 2 μl was used per qPCR reaction. To complete the reaction volume, 0.67 mM primers and SYBR^®^ Select Master Mix (Life Technologies) were added. The primers used are listed in Table 2.

**Table 2.**
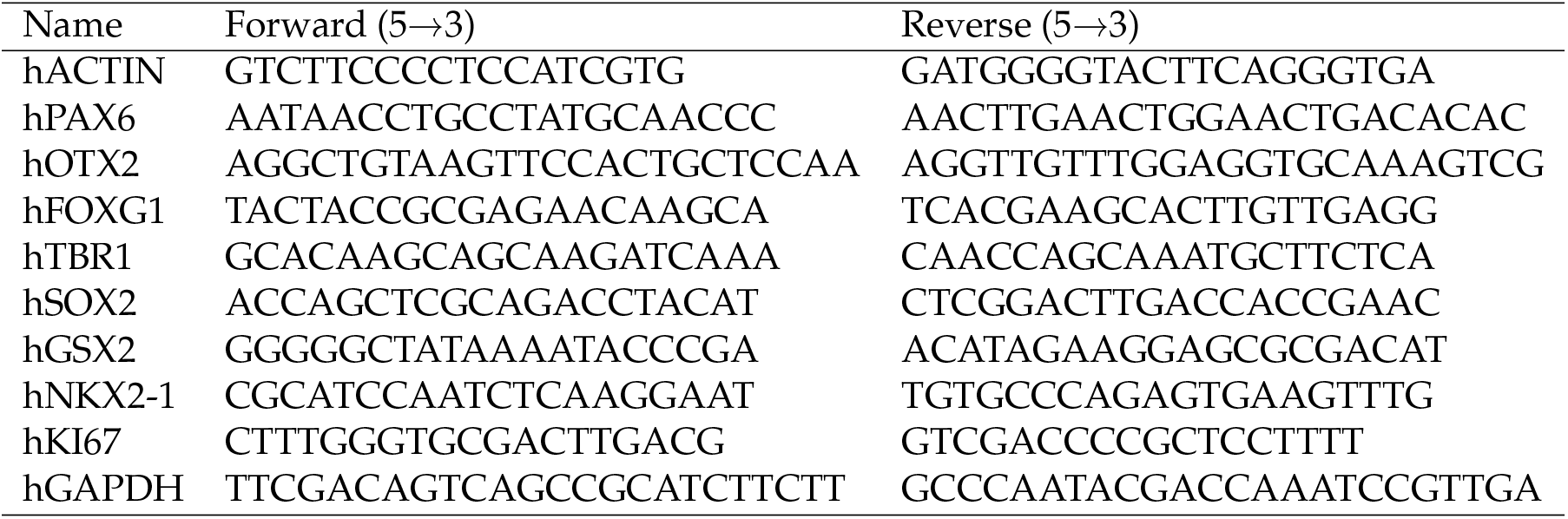
Sequences of primers used in qRT-PCR analysis.

### RNA-Seq and transcriptional analysis

RNA was extracted from organoids using an RNeasy mini kit (Qiagen) according to the manufacturer’s instructions. Subsequent library preparation (NEBNext^®^ Ultra™ RNA Library Prep Kit for Illumina) and paired end 2× 150 bp sequencing using HiSeq were performed by Novogene, Hong Kong. Data quality was assessed using FastQC [62]. The reads were aligned to the hg38 genome (Ensembl version 79) using STAR 2.5.2a [63] and quantified using RSEM 1.2.30 [64]. Mitochondrial and ribosomal genes were removed from further analysis to avoid confounding technical factors (e.g., differences in mitochondrial numbers and rRNA abundance). Differential expression was calculated using the Wald test within DESeq2 [65] using tximport [66]. For genes expressed over one transcript per million, differences in expression levels were deemed significant at a BH-corrected 10% false-discovery rate (FDR) level. Fold changes were calculated for IsoHD-81Q with respect to IsoHD-30Q lines. Functional enrichment and de novo motif discovery were performed with HOMER [37] using the list of differentially expressed genes in IsoHD-81Q versus IsoHD-30Q.

### Correlation of transcriptome between organoids and human brain development

The published RNA-seq data of three cortical sub-regions (dorsolateral prefrontal cortex, ventrolateral prefrontal cortex, and orbital frontal cortex) at nine developmental stages (8,12, 13, 16, 17, 21, 24, and 37 pcw and 4 months) of human brains were obtained from Allen Brain Atlas (http://www.brainspan.org/static/download.html). This dataset was compared with the organoids RNA-seq data of the current study to assess the degree of transcriptome correlation. For each cortical sub-region, pairwise Spearman correlation coefficients were computed between the different developmental stages of human brains and the IsoHD-30Q and IsoHD-81Q organoids using log normalized gene expression values. The correlation matrices were then colored on the same scale and visualized using R.

### FUCCI cell cycle analysis

Lentivirus FUCCI construct pBOB-EF1-FastFUCCI-Puro was a gift from Kevin Brindle Duncan Jodrell (Addgene plasmid 86849). For virus production, lentiviral vectors were co-transfected with packaging vectors into 293T cells and the supernatant was harvested and filtered through a 0.45-μm low protein binding cellulose acetate filter. Titers were then determined by infecting 293T cells with a serial dilution of the virus. For a typical preparation, the titer was approximately 1–5×10^6^/ml. hESC-derived neural progenitor cells (2×10^5^) were incubated in suspension with 1×10^6^ viral particles and 8 μg/ml polybrene for 3 h in a 37 °C incubator. The cells were then re-plated in 24-well plates and imaged with live-cell imaging Olympus IX-83.

### ATM inhibitor treatment

Organoids were treated with the ATM inhibitor KU60019 (5 μM) from day 14 to day 21 of differentiation. Organoids continued to grow to day 28 after treatment withdrawal. Six organoids for each line were sectioned at 200mm intervals and 5 continuous sections were analyzed. Radially organized tissues greater than 400μm or between 200μm and 400μm were counted for each line.

### Western blot

Cells were lysed on ice in RIPA buffer supplemented with protease inhibitor cocktail (Roche). Cell lysates were then sonicated (Qsonica-Q700 Sonicator, Misonix), and clarified by centrifugation at 14,000 rpm at 4 °C. Protein extracts were quantified using the Pierce BCA protein assay kit (Thermo Scientific). Equal amounts of proteins were resolved by SDS-PAGE and transferred onto nitrocellulose membranes. Membranes were probed with the listed primary and secondary antibodies, and developed using ECL on X-ray film.

### Statistical analysis

Unless otherwise stated, data are presented as mean ± standard error of the mean (SEM). Comparisons between groups were assessed using a one-way ANOVA with Fisher’s LSD post hoc analysis. Where indicated, pair-wise comparisons were assessed with Mann-Whitney U test or Student’s t-test. GraphPad Prism v7 software was used to analyse data for statistical significance. Differences were considered statistically significant when p < 0.05.

## Acknowledgments

We thank Amanda Chern for technical assistance. The work was partly funded by a Strategic Positioning Fund for Genetic Orphan Diseases (SPF2012/005) from the Agency for Science Technology and Research (Singapore), and an Industry Alignment Fund (SUREKids) to M.A.P. We thank the NINDS iPSC Repository for the HD hiPSC lines (18Q, 71Q, and 109Q).

## Author contributions

J.Z., J.O., K.H.U., and O.A.A. performed experiments; J.Z., O.A.A., S.R.L., S.M., and M.A.P. analyzed data; F.G. provided reagents; M.R., J.K., F.G., and E.P. provided technical advice and intellectual guidance; M.A.P. and J.Z. conceptualized the study, designed research, and wrote the paper.

---

**Figure S1.**
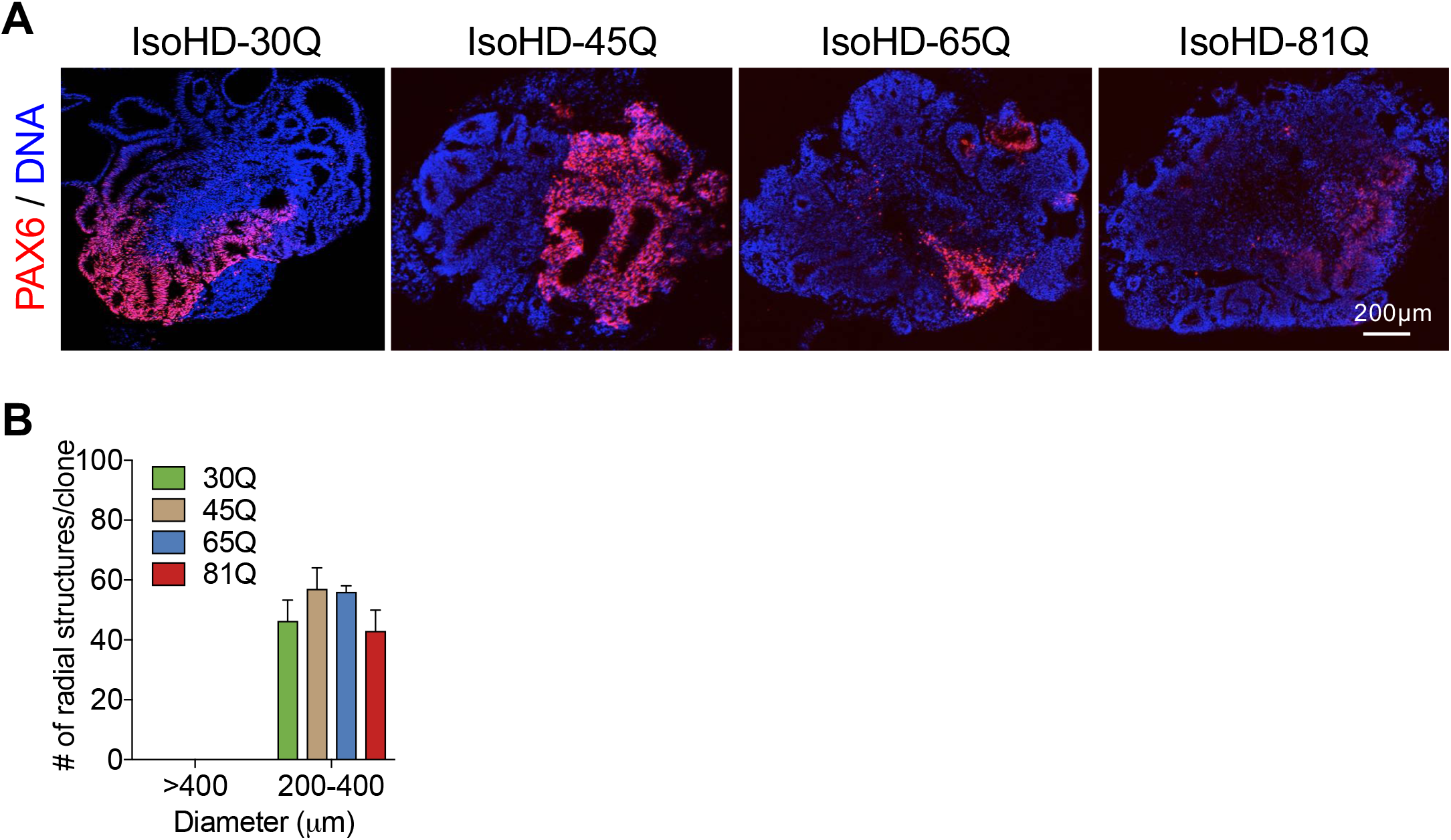
No difference in size of neuroepithelial structures in IsoHD hPSC-derived cerebral organoids at day 21. **(A)** Immunostaining of organoid sections. Representative images of IsoHD-30Q, 45Q, 65Q and 81Q organoids in low magnification showing PAX6-positive radial (neuroepithelial) structures. Scale bar: 200μm. **(B)** Analysis of organoids at day 21 of differentiation shows no difference in the size of radially-organized PAX6-positive, neuroepithelial structures between the different IsoHD lines. 6 organoids per IsoHD line were analyzed. Values represent mean±SEM.

**Figure S2.**
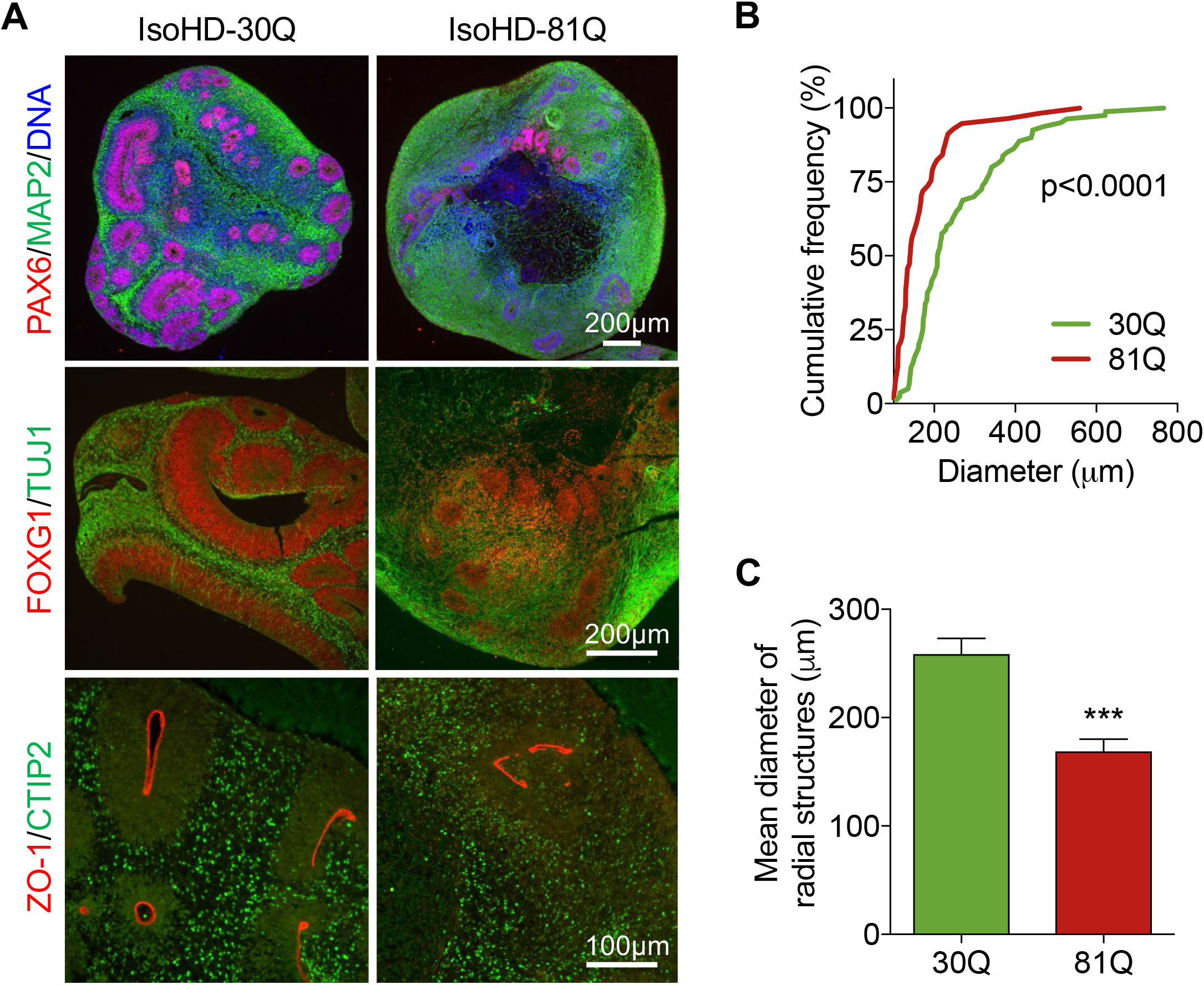
Reduced size of neuroepithelial structures in IsoHD-81Q hPSC-derived cerebral organoids at day 42. **(A)** Immunostaining of organoid sections. Representative images of IsoHD-30Q and 81Q organoids showing PAX6-positive radial (neuroepithelial) structures, expression of FOXG1 forebrain and CTIP2 cortical markers. **(B)** Cumulative frequency analysis showing a shift towards smaller radial structures in IsoHD-81Q organoids compared with IsoHD-30Q (p<0.0001 by the unpaired two-tailed Kolmogorov-Smirnov test). **(C)** Mean diameter of radial neuroepithelial structures in IsoHD-30Q and IsoHD-81Q cerebral organoids. 57-80 structures from 6 organoids per IsoHD line were analyzed. Values represent mean±SEM.

**Figure S3.**
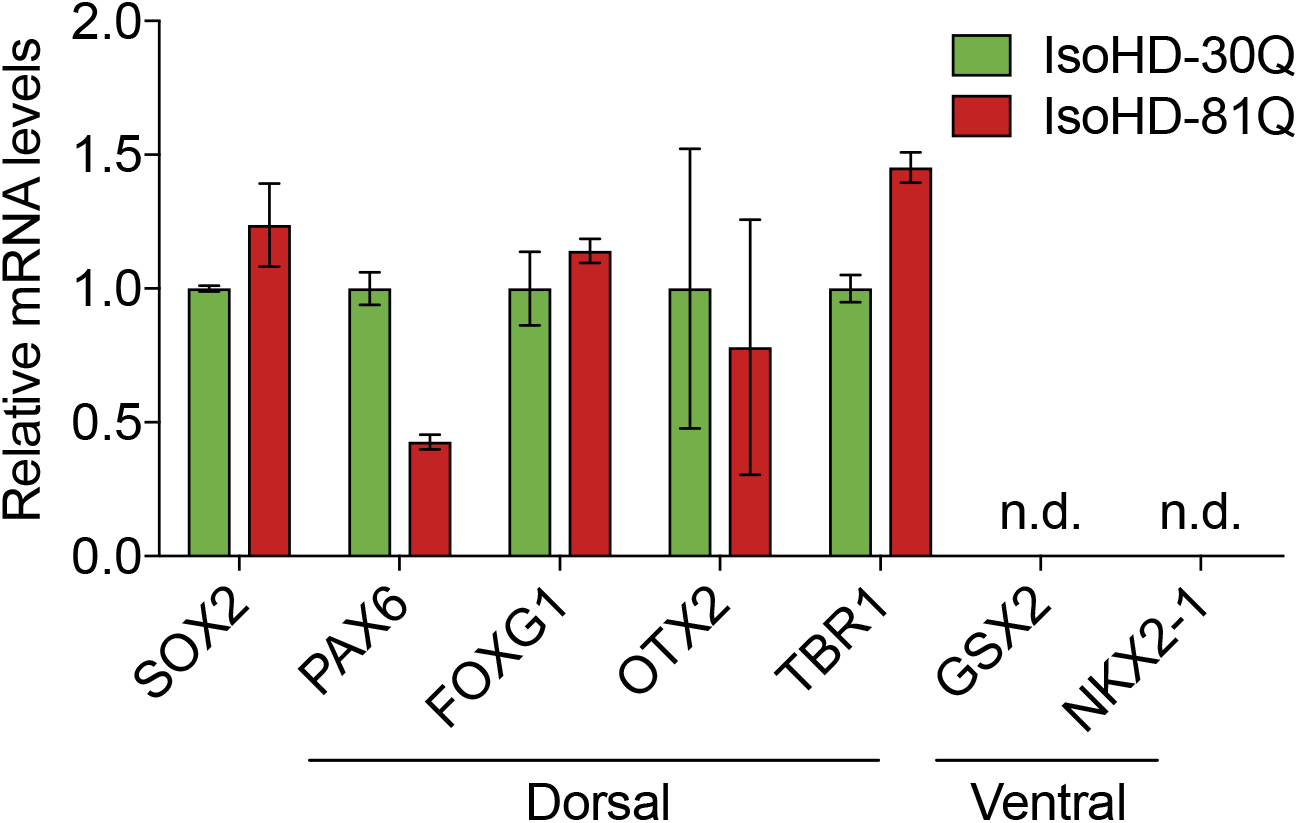
Expression of dorsal forebrain markers by IsoHD cerebral organoids. Dorsal and ventral forebrain marker analysis by qRT-PCR in day 28 IsoHD organoids, (n=3/genotype). Values represent mean±SEM.

**Figure S4.**
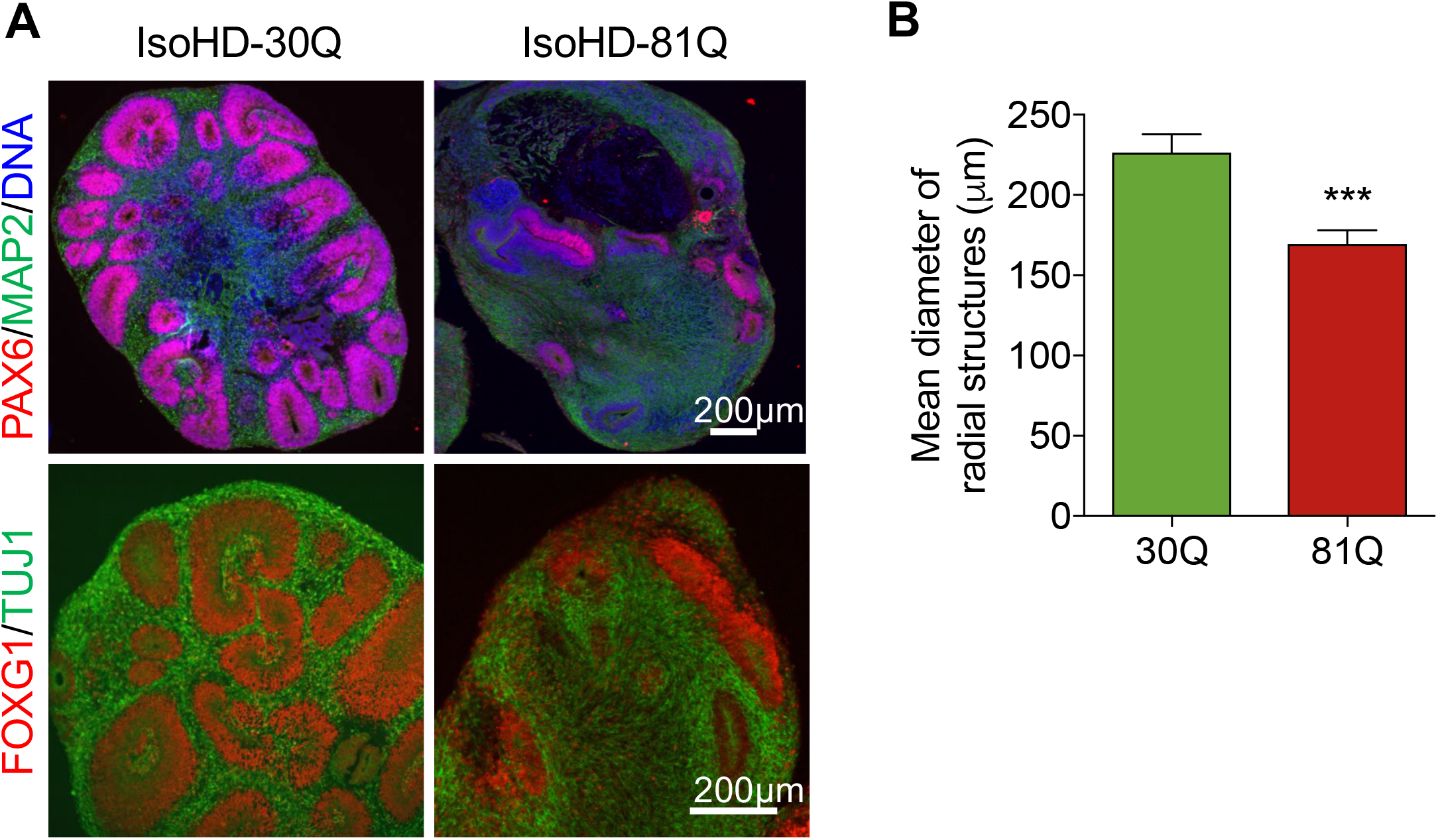
The reduction in the size of neuroepithelial structures is maintained following forebrain-specific patterning in IsoHD-81Q hPSC-derived cerebral organoids at day 28. **(A)** Immunostaining of organoid sections. Representative images of IsoHD-30Q and 81Q organoids showing PAX6-positive radial (neuroepithelial) structures and expression of FOXG1 forebrain marker. **(B)** Mean diameter of radial neuroepithelial structures in IsoHD-30Q and IsoHD-81Q cerebral organoids. 67-77 structures from 6 organoids per IsoHD line were analyzed. Values represent mean±SEM.

**Figure S5.**
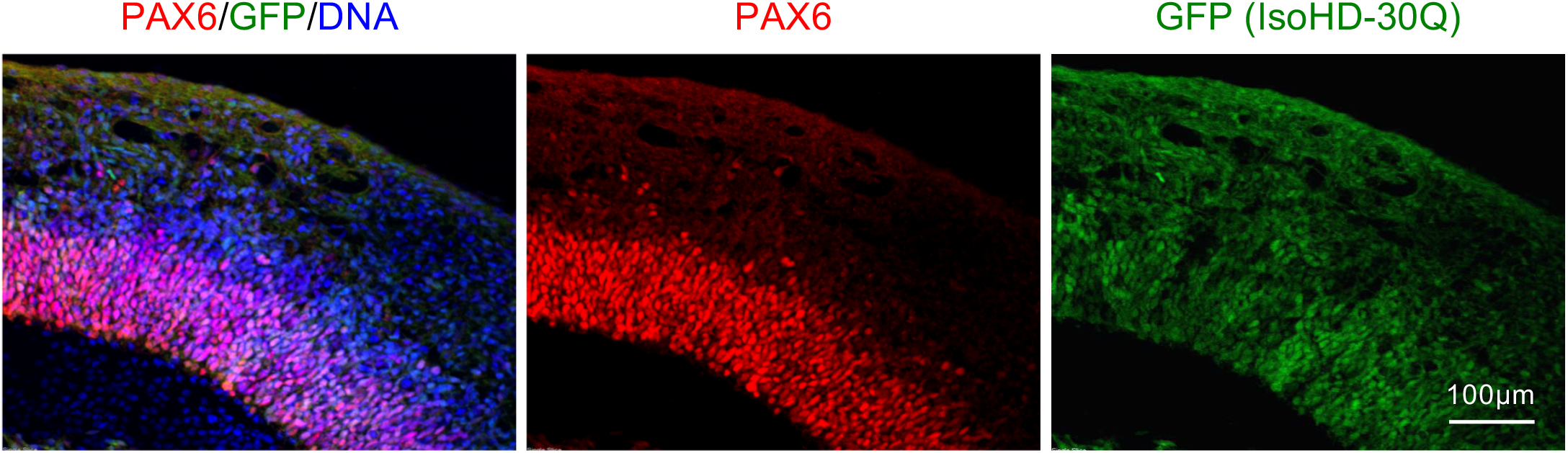
Neuroepithelial development in GFP-labeled IsoHD-30Q organoids. Neuroepithelial development in IsoHD-30Q/30Q-GFP organoids at day 28. Organoid sections were stained for GFP and PAX6.

## References

[1] G. P. Bates, R. Dorsey, J. F. Gusella, M. R. Hayden, C. Kay, B. R. Leavitt, M. Nance, C. A. Ross, R. I. Scahill, R. Wetzel, E. J. Wild, and S. J. Tabrizi, “Huntington disease,” Nature Reviews Disease Primers, vol. 1, p. 15005, Dec. 2015.

[2] M. A. Nance and R. H. Myers, “Juvenile onset Huntington’s disease?clinical and research perspectives,” Mental Retardation and Developmental Disabilities Research Reviews, vol. 7, pp. 153–157, Aug. 2001.

[3] P. C. Nopoulos, E. H. Aylward, C. A. Ross, J. A. Mills, D. R. Langbehn, H. J. Johnson, V. A. Magnotta, R. K. Pierson, L. J. Beglinger, M. A. Nance, R. A. Barker, J. S. Paulsen, and the PREDICT-HD Investigators and Coordinators of the Huntington Study Group, “Smaller intracranial volume in prodromal Huntington’s disease: evidence for abnormal neurodevelopment,” Brain, vol. 134, pp. 137–142, Jan. 2011.

[4] J. K. Lee, K. Mathews, B. Schlaggar, J. Perlmutter, J. S. Paulsen, E. Epping, L. Burmeister, and P. Nopoulos, “Measures of growth in children at risk for Huntington disease,” Neurology, vol. 79, pp. 668–674, Aug. 2012.

[5] J. S. Paulsen, V. A. Magnotta, A. E. Mikos, H. L. Paulson, E. Penziner, N. C. Andreasen, and P. C. Nopoulos, “Brain Structure in Preclinical Huntington’s Disease,” Biological Psychiatry, vol. 59, pp. 57–63, Jan. 2006.

[6] E. van der Plas, D. R. Langbehn, A. L. Conrad, T. R. Koscik, A. Tereshchenko, E. A. Epping, V. A. Magnotta, and P. C. Nopoulos, “Abnormal brain development in child and adolescent carriers of mutant huntingtin,” Neurology, vol. 93, pp. e1021–e1030, Sept. 2019.

[7] F. Lopes, M. Barbosa, A. Ameur, G. Soares, J. de Sa, A. I. Dias, G. Oliveira, P. Cabral, T. Temudo, E. Calado, I. F. Cruz, J. P. Vieira, R. Oliveira, S. Esteves, S. Sauer, I. Jonasson, A.-C. Syvanen, U. Gyllensten, D. Pinto, and P. Maciel, “Identification of novel genetic causes of Rett syndrome-*like* phenotypes,” Journal of Medical Genetics, vol. 53, pp. 190–199, Mar. 2016.

[8] L. H. Rodan, J. Cohen, A. Fatemi, T. Gillis, D. Lucente, J. Gusella, and J. D. Picker, “A novel neurodevelopmental disorder associated with compound heterozygous variants in the huntingtin gene,” European Journal of Human Genetics, vol. 24, pp. 1826–1827, Dec. 2016.

[9] M. F. Mehler, J. R. Petronglo, E. E. Arteaga-Bracho, M. E. Gulinello, M. L. Winchester, N. Pichamoorthy, S. K. Young, C. D. DeJesus, H. Ishtiaq, S. Gokhan, and A. E. Molero, “Loss-of-Huntingtin in Medial and Lateral Ganglionic Lineages Differentially Disrupts Regional Interneuron and Projection Neuron Subtypes and Promotes Huntington’s Disease-Associated Behavioral, Cellular, and Pathological Hallmarks,” The Journal of Neuroscience, vol. 39, pp. 1892–1909, Mar. 2019.

[10] A. Reiner, N. Del Mar, C. A. Meade, H. Yang, I. Dragatsis, S. Zeitlin, and D. Goldowitz, “Neurons Lacking Huntingtin Differentially Colonize Brain and Survive in Chimeric Mice,” The Journal of Neuroscience, vol. 21, pp. 7608–7619, Oct. 2001.

[11] J. K. White, W. Auerbach, M. P. Duyao, J.-P. Vonsattel, J. F. Gusella, A. L. Joyner, and M. E. MacDonald, “Huntingtin is required for neurogenesis and is not impaired by the Huntington’s disease CAG expansion,” Nature Genetics, vol. 17, pp. 404–410, Dec. 1997.

[12] J. D. Godin, K. Colombo, M. Molina-Calavita, G. Keryer, D. Zala, B. C. Charrin, P. Dietrich, M.-L. Volvert, F. Guillemot, I. Dragatsis, Y. Bellaiche, F. Saudou, L. Nguyen, and S. Humbert, “Huntingtin Is Required for Mitotic Spindle Orientation and Mammalian Neurogenesis,” Neuron, vol. 67, pp. 392–406, Aug. 2010.

[13] G. D. Nguyen, S. Gokhan, A. E. Molero, and M. F. Mehler, “Selective Roles of Normal and Mutant Huntingtin in Neural Induction and Early Neurogenesis,” PLoS ONE, vol. 8, p. e64368, May 2013.

[14] M. Barnat, J. Le Friec, C. Benstaali, and S. Humbert, “Huntingtin-Mediated Multipolar-Bipolar Transition of Newborn Cortical Neurons Is Critical for Their Postnatal Neuronal Morphology,” Neuron, vol. 93, pp. 99–114, Jan. 2017.

[15] Y. Tong, T. J. Ha, L. Liu, A. Nishimoto, A. Reiner, and D. Goldowitz, “Spatial and Temporal Requirements for huntingtin (Htt) in Neuronal Migration and Survival during Brain Development,” Journal of Neuroscience, vol. 31, pp. 14794–14799, Oct. 2011.

[16] J. A. Cooper, “Molecules and mechanisms that regulate multipolar migration in the intermediate zone,” Frontiers in Cellular Neuroscience, vol. 8, Nov. 2014.

[17] O. Marin and J. L. Rubenstein, “Cell migration in the forebrain,” Annual Review of Neuroscience, vol. 26, pp. 441–483, Mar. 2003.

[18] C. Lopes, S. Aubert, F. Bourgois-Rocha, M. Barnat, A. C. Rego, N. Deglon, A. L. Perrier, and S. Humbert, “Dominant-Negative Effects of Adult-Onset Huntingtin Mutations Alter the Division of Human Embryonic Stem Cells-Derived Neural Cells,” PLOS ONE, vol. 11, p. e0148680, Feb. 2016.

[19] A. Ruzo, G. F. Croft, J. J. Metzger, S. Galgoczi, L. J. Gerber, C. Pellegrini, H. Wang, M. Fenner, S. Tse, A. Marks, C. Nchako, and A. H. Brivanlou, “Chromosomal instability during neurogenesis in Huntington’s disease,” Development, vol. 145, p. dev156844, Jan. 2018.

[20] M. Molina-Calavita, M. Barnat, S. Elias, E. Aparicio, M. Piel, and S. Humbert, “Mutant Huntingtin Affects Cortical Progenitor Cell Division and Development of the Mouse Neocortex,” Journal of Neuroscience, vol. 34, pp. 10034–10040, July 2014.

[21] P. Conforti, S. Camnasio, C. Mutti, M. Valenza, M. Thompson, E. Fossale, S. Zeitlin, M. E. MacDonald, C. Zuccato, and E. Cattaneo, “Lack of huntingtin promotes neural stem cells differentiation into glial cells while neurons expressing huntingtin with expanded polyglutamine tracts undergo cell death,” Neurobiology of Disease, vol. 50, pp. 160–170, Feb. 2013.

[22] M. T. Lorincz and V. A. Zawistowski, “Expanded CAG repeats in the murine Huntington’s disease gene increases neuronal differentiation of embryonic and neural stem cells,” Molecular and Cellular Neuroscience, vol. 40, pp. 1–13, Jan. 2009.

[23] P. P. Mathkar, D. Suresh, J. Dunn, C. M. Tom, and V. B. Mattis, “Characterization of Neurodevelopmental Abnormalities in iPSC-Derived Striatal Cultures from Patients with Huntington’s Disease,” Journal of Huntington’s Disease, vol. 8, pp. 257–269, Aug. 2019.

[24] S. R. Mehta, C. M. Tom, Y. Wang, C. Bresee, D. Rushton, P. P. Mathkar, J. Tang, and V. B. Mattis, “Human Huntington’s Disease iPSC-Derived Cortical Neurons Display Altered Transcriptomics, Morphology, and Maturation,” Cell Reports, vol. 25, pp. 1081–1096.e6, Oct. 2018.

[25] K. L. Ring, M. C. An, N. Zhang, R. N. O’Brien, E. M. Ramos, F. Gao, R. Atwood, B. J. Bailus, S. Melov, S. D. Mooney, G. Coppola, and L. M. Ellerby, “Genomic Analysis Reveals Disruption of Striatal Neuronal Development and Therapeutic Targets in Human Huntington’s Disease Neural Stem Cells,” Stem Cell Reports, vol. 5, pp. 1023–1038, Dec. 2015.

[26] K. Świtońska, W. J. Szlachcic, L. Handschuh, P. Wojciechowski, Ł. Marczak, M. Stelmaszczuk, M. Figlerowicz, and M. Figiel, “Identification of Altered Developmental Pathways in Human Juvenile HD iPSC With 71q and 109q Using Transcriptome Profiling,” Frontiers in Cellular Neuroscience, vol. 12, p. 528, Jan. 2019.

[27] K. Wiatr, W. J. Szlachcic, M. Trzeciak, M. Figlerowicz, and M. Figiel, “Huntington Disease as a Neurodevelopmental Disorder and Early Signs of the Disease in Stem Cells,” Molecular Neurobiology, vol. 55, pp. 3351–3371, Apr. 2018.

[28] T. Haremaki, J. J. Metzger, T. Rito, M. Z. Ozair, F. Etoc, and A. H. Brivanlou, “Self-organizing neuru-loids model developmental aspects of Huntington’s disease in the ectodermal compartment,” Nature Biotechnology, vol. 37, pp. 1198–1208, Oct. 2019.

[29] P. Conforti, D. Besusso, V. D. Bocchi, A. Faedo, E. Cesana, G. Rossetti, V. Ranzani, C. N. Svendsen, L. M. Thompson, M. Toselli, G. Biella, M. Pagani, and E. Cattaneo, “Faulty neuronal determination and cell polarization are reverted by modulating HD early phenotypes,” Proceedings of the National Academy of Sciences, vol. 115, pp. E762–E771, Jan. 2018.

[30] M. A. Lancaster, M. Renner, C.-A. Martin, D. Wenzel, L. S. Bicknell, M. E. Hurles, T. Homfray, J. M. Penninger, A. P. Jackson, and J. A. Knoblich, “Cerebral organoids model human brain development and microcephaly,” Nature, vol. 501, pp. 373–379, Sept. 2013.

[31] J. Ooi, S. R. Langley, X. Xu, K. H. Utami, B. Sim, Y. Huang, N. P. Harmston, Y. L. Tay, A. Ziaei, R. Zeng, D. Low, F. Aminkeng, R. M. Sobota, F. Ginhoux, E. Petretto, and M. A. Pouladi, “Unbiased Profiling of Isogenic Huntington Disease hPSC-Derived CNS and Peripheral Cells Reveals Strong Cell-Type Specificity of CAG Length Effects,” Cell Reports, vol. 26, pp. 2494–2508.e7, Feb. 2019.

[32] I. Kelava and M. A. Lancaster, “Dishing out mini-brains: Current progress and future prospects in brain organoid research,” Developmental Biology, vol. 420, pp. 199–209, Dec. 2016.

[33] E. Taverna, M. Gotz, and W. B. Huttner, “The Cell Biology of Neurogenesis: Toward an Understanding of the Development and Evolution of the Neocortex,” Annual Review of Cell and Developmental Biology, vol. 30, pp. 465–502, Oct. 2014.

[34] M. A. Lancaster, N. S. Corsini, S. Wolfinger, E. H. Gustafson, A. W. Phillips, T. R. Burkard, T. Otani, F. J. Livesey, and J. A. Knoblich, “Guided self-organization and cortical plate formation in human brain organoids,” Nature Biotechnology, vol. 35, pp. 659–666, July 2017.

[35] X. Qian, H. N. Nguyen, M. M. Song, C. Hadiono, S. C. Ogden, C. Hammack, B. Yao, G. R. Hamersky, F. Jacob, C. Zhong, K.-j. Yoon, W. Jeang, L. Lin, Y. Li, J. Thakor, D. A. Berg, C. Zhang, E. Kang, M. Chickering, D. Nauen, C.-Y. Ho, Z. Wen, K. M. Christian, P.-Y. Shi, B. J. Maher, H. Wu, P. Jin, H. Tang, H. Song, and G.-l. Ming, “Brain-Region-Specific Organoids Using Mini-bioreactors for Modeling ZIKV Exposure,” Cell, vol. 165, pp. 1238–1254, May 2016.

[36] The HD iPSC Consortium, “Developmental alterations in Huntington’s disease neural cells and pharmacological rescue in cells and mice,” Nature Neuroscience, vol. 20, pp. 648–660, May 2017.

[37] S. Heinz, C. Benner, N. Spann, E. Bertolino, Y. C. Lin, P. Laslo, J. X. Cheng, C. Murre, H. Singh, and C. K. Glass, “Simple Combinations of Lineage-Determining Transcription Factors Prime cis-Regulatory Elements Required for Macrophage and B Cell Identities,” Molecular Cell, vol. 38, pp. 576–589, May 2010.

[38] C. Bertoli, J. M. Skotheim, and R. A. M. de Bruin, “Control of cell cycle transcription during G1 and S phases,” Nature Reviews Molecular Cell Biology, vol. 14, pp. 518–528, Aug. 2013.

[39] J. Musa, M.-M. Aynaud, O. Mirabeau, O. Delattre, and T. G. Grünewald, “MYBL2 (B-Myb): a central regulator of cell proliferation, cell survival and differentiation involved in tumorigenesis,” Cell Death & Disease, vol. 8, pp. e2895–e2895, June 2017.

[40] C. F. Schaefer, K. Anthony, S. Krupa, J. Buchoff, M. Day, T. Hannay, and K. H. Buetow, “PID: the Pathway Interaction Database,” Nucleic Acids Research, vol. 37, pp. D674–D679, Jan. 2009.

[41] C. Dehay and H. Kennedy, “Cell-cycle control and cortical development,” Nature Reviews Neuroscience, vol. 8, pp. 438–450, June 2007.

[42] P. Salomoni and F. Calegari, “Cell cycle control of mammalian neural stem cells: putting a speed limit on G1,” Trends in Cell Biology, vol. 20, pp. 233–243, May 2010.

[43] T. Takahashi, R. Nowakowski, and V. Caviness, “The cell cycle of the pseudostratified ventricular epithelium of the embryonic murine cerebral wall,” The Journal of Neuroscience, vol. 15, pp. 6046–6057, Sept. 1995.

[44] F. Calegari, “An inhibition of cyclin-dependent kinases that lengthens, but does not arrest, neuroepithelial cell cycle induces premature neurogenesis,” Journal of Cell Science, vol. 116, pp. 4947–4955, Dec. 2003.

[45] C. Lange, W. B. Huttner, and F. Calegari, “Cdk4/CyclinD1 Overexpression in Neural Stem Cells Shortens G1, Delays Neurogenesis, and Promotes the Generation and Expansion of Basal Progenitors,” Cell Stem Cell, vol. 5, pp. 320–331, Sept. 2009.

[46] A. Sakaue-Sawano, H. Kurokawa, T. Morimura, A. Hanyu, H. Hama, H. Osawa, S. Kashiwagi, K. Fukami, T. Miyata, H. Miyoshi, T. Imamura, M. Ogawa, H. Masai, and A. Miyawaki, “Visualizing Spatiotemporal Dynamics of Multicellular Cell-Cycle Progression,” Cell, vol. 132, pp. 487–498, Feb. 2008.

[47] A. Shitamukai and F. Matsuzaki, “Control of asymmetric cell division of mammalian neural progenitors,” Development, Growth & Differentiation, vol. 54, pp. 277–286, Apr. 2012.

[48] M. F. Lavin, “Ataxia-telangiectasia: from a rare disorder to a paradigm for cell signalling and cancer,” Nature Reviews Molecular Cell Biology, vol. 9, pp. 759–769, Oct. 2008.

[49] X.-H. Lu, V. B. Mattis, N. Wang, I. Al-Ramahi, N. van den Berg, S. A. Fratantoni, H. Waldvogel, E. Greiner, A. Osmand, K. Elzein, J. Xiao, S. Dijkstra, R. de Pril, H. V. Vinters, R. Faull, E. Signer, S. Kwak, J. J. Marugan, J. Botas, D. F. Fischer, C. N. Svendsen, I. Munoz-Sanjuan, and X. W. Yang, “Targeting ATM ameliorates mutant Huntingtin toxicity in cell and animal models of Huntington’s disease,” Science Translational Medicine, vol. 6, pp. 268ra178–268ra178, Dec. 2014.

[50] S. E. Golding, E. Rosenberg, N. Valerie, I. Hussaini, M. Frigerio, X. F. Cockcroft, W. Y. Chong, M. Hummer-sone, L. Rigoreau, K. A. Menear, M. J. O’Connor, L. F. Povirk, T. van Meter, and K. Valerie, “Improved ATM kinase inhibitor KU-60019 radiosensitizes glioma cells, compromises insulin, AKT and ERK prosurvival signaling, and inhibits migration and invasion,” Molecular Cancer Therapeutics, vol. 8, pp. 2894–2902, Oct. 2009.

[51] T. Namba and W. B. Huttner, “Neural progenitor cells and their role in the development and evolutionary expansion of the neocortex: Neural progenitor cells’ role in the development and evolutionary expansion of the neocortex,” Wiley Interdisciplinary Reviews: Developmental Biology, vol. 6, p. e256, Jan. 2017.

[52] F. Pinto-Teixeira and C. Desplan, “Notch activity in neural progenitors coordinates cytokinesis and asymmetric differentiation,” Science Signaling, vol. 7, pp. pe26–pe26, Oct. 2014.

[53] D. M. Allen, “Ataxia telangiectasia mutated is essential during adult neurogenesis,” Genes & Development, vol. 15, pp. 554–566, Mar. 2001.

[54] J.-I. Chae, D.-W. Kim, N. Lee, Y.-J. Jeon, I. Jeon, J. Kwon, J. Kim, Y. Soh, D.-S. Lee, K. S. Seo, N.-J. Choi, B. C. Park, S. H. Kang, J. Ryu, S.-H. Oh, D. A. Shin, D. R. Lee, J. T. Do, I.-H. Park, G. Q. Daley, and J. Song, “Quantitative proteomic analysis of induced pluripotent stem cells derived from a human Huntington’s disease patient,” Biochemical Journal, vol. 446, pp. 359–371, Sept. 2012.

[55] P. Giuliano, “DNA damage induced by polyglutamine-expanded proteins,” Human Molecular Genetics, vol. 12, pp. 2301–2309, July 2003.

[56] J. Illuzzi, S. Yerkes, H. Parekh-Olmedo, and E. B. Kmiec, “DNA breakage and induction of DNA damage response proteins precede the appearance of visible mutant huntingtin aggregates,” Journal of Neuroscience Research, vol. 87, pp. 733–747, Feb. 2009.

[57] J. H. Sun, L. Zhou, D. J. Emerson, S. A. Phyo, K. R. Titus, W. Gong, T. G. Gilgenast, J. A. Beagan, B. L. Davidson, F. Tassone, and J. E. Phillips-Cremins, “Disease-Associated Short Tandem Repeats Co-localize with Chromatin Domain Boundaries,” Cell, vol. 175, pp. 224–238.e15, Sept. 2018.

[58] A. E. Molero, E. E. Arteaga-Bracho, C. H. Chen, M. Gulinello, M. L. Winchester, N. Pichamoorthy, S. Gokhan, K. Khodakhah, and M. F. Mehler, “Selective expression of mutant huntingtin during development recapitulates characteristic features of Huntington’s disease,” Proceedings of the National Academy of Sciences, vol. 113, pp. 5736–5741, May 2016.

[59] C. Zuccato and E. Cattaneo, “HTT Evolution and Brain Development,” in Programmed Cells from Basic Neuroscience to Therapy, pp. 41–55, Berlin, Heidelberg: Springer Berlin Heidelberg, Apr. 2013.

[60] M. Tartari, C. Gissi, V. Lo Sardo, C. Zuccato, E. Picardi, G. Pesole, and E. Cattaneo, “Phylogenetic Comparison of Huntingtin Homologues Reveals the Appearance of a Primitive polyQ in Sea Urchin,” Molecular Biology and Evolution, vol. 25, pp. 330–338, Jan. 2008.

[61] V. B. Mattis, C. Tom, S. Akimov, J. Saeedian, M. E. Østergaard, A. L. Southwell, C. N. Doty, L. Ornelas, A. Sahabian, L. Lenaeus, B. Mandefro, D. Sareen, J. Arjomand, M. R. Hayden, C. A. Ross, and C. N. Svendsen, “HD iPSC-derived neural progenitors accumulate in culture and are susceptible to BDNF withdrawal due to glutamate toxicity,” Human Molecular Genetics, vol. 24, pp. 3257–3271, June 2015.

[62] S. Andrews, “FastQC: a quality control tool for high throughput sequence data,” 2010.

[63] A. Dobin, C. A. Davis, F. Schlesinger, J. Drenkow, C. Zaleski, S. Jha, P. Batut, M. Chaisson, and T. R. Gingeras, “STAR: ultrafast universal RNA-seq aligner,” Bioinformatics, vol. 29, pp. 15–21, Jan. 2013.

[64] B. Li and C. N. Dewey, “RSEM: accurate transcript quantification from RNA-Seq data with or without a reference genome,” BMC Bioinformatics, vol. 12, p. 323, Dec. 2011.

[65] M. I. Love, W. Huber, and S. Anders, “Moderated estimation of fold change and dispersion for RNA-seq data with DESeq2,” Genome Biology, vol. 15, p. 550, Dec. 2014.

[66] C. Soneson, M. I. Love, and M. D. Robinson, “Differential analyses for RNA-seq: transcript-level estimates improve gene-level inferences,” F1000Research, vol. 4, p. 1521, Feb. 2016.

